# APOBEC3-driven neoantigen-rich cancers co-opt 1q23.3 amplification for tumor-intrinsic immune cloaking

**DOI:** 10.64898/2026.07.01.735125

**Authors:** Dhanusha Yesudhas, Bilal Lone, Ecem Unal, Arup Chakraborty, Ayse G. Keskus, Jaimie Ryou, Kelly Butler, Tania Colon Aquino, Abbas Yousefi-Rad, Weiming Yang, Lisa M Jenkins, Raju Chelluri, Elias BA Chandran, Vladimir A. Valera Romero, Salah Boudjadi, Sandeep Gurram, Mikhail Kolmogorov, Andrea B. Apolo, A. Rouf Banday

## Abstract

Hypermutational processes, including those driven by the APOBEC3 family of cytidine deaminases, generate abundant neoantigens yet give rise to tumors that evade immune recognition. Here, using multi-omics analyses followed by functional validation, we identified a tumor-intrinsic immune-cloaking mechanism in neoantigen-rich epithelial cancers, characterized by coordinated suppression of antigen presentation, immune-recruiting cytokines and immune-checkpoint programs. In bladder cancer, genome-wide copy-number analysis identified recurrent 1q23.3 amplification as a genomic feature of a neoantigen-high/CD8-low tumor state. Within this locus, *NECTIN4* emerged as the dominant candidate effector, outperforming extrachromosomal DNA status as a predictor of immune–neoantigen discordance. Similar associations were observed across breast and lung cancers. Functional studies demonstrated that NECTIN4 was sufficient to establish a T-cell-poor tumor microenvironment and confer resistance to PD-1 blockade in immunocompetent mice. Mechanistically, NECTIN4 engaged a DDR1–SHP2 axis that suppressed STAT1 phosphorylation, silencing tumor-cell immune-engagement programs. NECTIN4 blockade restored STAT1 activity and reduced tumor growth, indicating that the cloaked state is pharmacologically reversible. Mutational signature, breakpoint motif, timing and clonality analyses, together with *APOBEC3B* expression and germline genetic evidence, linked APOBEC3-mediated mutagenesis to recurrent 1q23.3 amplification encompassing *NECTIN4*. These findings reveal how neoantigen-generating mutational processes can be coupled to structural genome evolution to enable tumor-intrinsic immune cloaking through a therapeutically targetable NECTIN4–DDR1–SHP2 axis.

## Main

Tumor immune surveillance imposes strong selective pressure on cancer evolution^1^, favoring mechanisms that prevent immune recognition or attenuate immune attack. This creates a paradox for hypermutated cancers driven by processes such as APOBEC3 cytidine deaminase activity, mismatch-repair deficiency and other mutagenic programs. Because elevated tumor mutational burden increases the probability of generating neoantigens, hypermutated tumors would be expected to promote immune engagement and establish CD8⁺ T cell–rich tumor microenvironments. Yet many such tumors remain immune-cold and T cell–poor^2–5^, indicating that neoantigen generation can be counterbalanced by tumor-intrinsic programs that suppress immune recognition despite high antigenic potential.

This paradox has direct implications for cancer immunotherapy^6,7^. Immune-checkpoint blockade has transformed outcomes across multiple malignancies, and tumor mutational burden is used as a biomarker of checkpoint responsiveness in part because it serves as a proxy for neoantigen abundance^8–11^. However, the predictive performance of tumor mutational burden varies substantially across tumor types and clinical cohorts, underscoring the need to define tumor-intrinsic mechanisms that uncouple neoantigen burden from productive antitumor immunity and that could be therapeutically targeted to improve immunotherapy response^12,13^.

APOBEC3-mediated mutagenesis is a major contributor to elevated mutational burden across epithelial malignancies, including bladder, lung and breast cancers^14–17^. These tumors are also enriched for copy-number alterations (CNAs)^18–20^, and complex genomic rearrangements, including extrachromosomal DNA (ecDNA), features increasingly linked to immune-suppressive tumor states^21–23^. However, how structural genome alterations shape immune visibility in neoantigen-rich, APOBEC3-mutagenized tumors remains poorly understood.

Here, we integrate large-scale tumor multi-omics and single-cell datasets with functional perturbation studies in vitro and in vivo to define a mechanism that decouples neoantigenicity from immune recognition in hypermutated epithelial cancers. We show that APOBEC3-driven tumors recurrently acquire chromosome 1q23.3 copy-number gains, with *NECTIN4* emerging as the predominant gene-level predictor of a neoantigen-high and CD8-low immune-discordant state. Functional studies identify NECTIN4 as a cell-surface regulator of STAT1-dependent immune-engagement programs, establishing a tumor-intrinsic suppressive state that we term tumor-intrinsic immune cloaking. These findings reveal a genome-encoded mechanism by which APOBEC3 mutagenesis, structural genome evolution and tumor-cell-intrinsic immune suppression converge to enable immune escape in neoantigen-rich cancers.

## Results

### Genome-wide analyses link copy-number alterations and ecDNA to immune–neoantigen discordance

To identify mechanisms of immune escape in tumors with high antigenic potential, we first analyzed The Cancer Genome Atlas bladder cancer cohort (TCGA-BLCA), a malignancy with frequent hypermutation and prominent APOBEC3-associated mutagenesis^14,17,24^. We performed a regression-based residual analysis relating predicted neoantigen burden to CD8⁺ T-cell infiltration inferred from bulk RNA sequencing, thereby identifying tumors that deviated from the expected positive relationship between antigenicity and immune infiltration. This approach defined a subset of tumors with high neoantigen burden but low CD8⁺ T-cell infiltration, hereafter referred to as NeoAg^hi^CD8^low^ tumors (n = 60; **Fig. 1a**). Classification of this subset was highly concordant after adjustment for tumor purity and ploidy (Cohen’s κ = 0.92; **Supplementary Fig. 1a**). As expected, these tumors displayed significantly elevated neoantigen load (*P* < 2.0 × 10⁻¹⁶) and markedly reduced CD8⁺ T-cell infiltration (*P* = 3.8 × 10⁻¹¹; **Fig. 1b**).

**Fig. 1.**
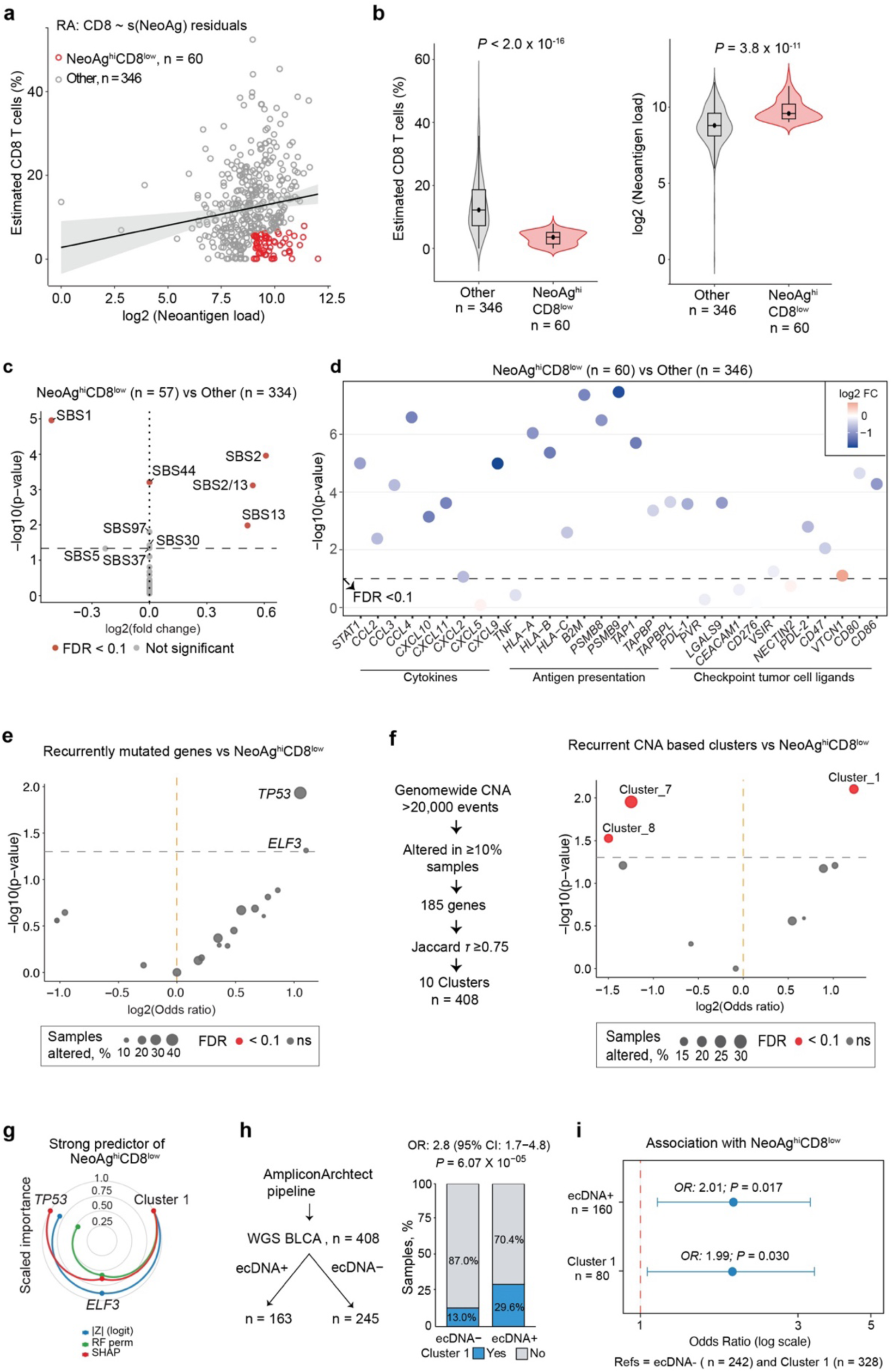
Genome-wide associations of CNAs and ecDNA status with the NeoAg^hi^CD8^low^ tumor state. a, Scatter plot of tumor neoantigen load versus estimated CD8⁺ T-cell abundance in TCGA bladder cancer (BLCA) tumors. CD8⁺ T-cell abundance was inferred from bulk RNA-seq deconvolution. NeoAg^hi^CD8^low^ tumors were identified using residual regression analysis of CD8⁺ T-cell abundance relative to neoantigen load. **b,** Comparison of estimated CD8⁺ T-cell abundance and neoantigen load between NeoAg^hi^CD8^low^ tumors and other BLCA tumors. *P* values were calculated using two-sided Mann–Whitney U tests. **c,** Volcano plot showing differential enrichment of mutational signatures in NeoAg^hi^CD8^low^ tumors compared with other BLCA tumors. The x axis shows log₂ fold change, and the y axis shows −log₁₀(*P* value). Red points indicate signatures significant at FDR < 0.1. **d,** Differential expression of cytokine, antigen-presentation and immune-checkpoint ligand gene panels between NeoAg^hi^CD8^low^ tumors and other BLCA tumors. The y axis shows −log₁₀(P value) from two-sided Wilcoxon rank-sum tests, and dot color indicates log₂ fold change in NeoAg^hi^CD8^low^ tumors relative to other tumors. The dashed line denotes FDR < 0.1. **e,** Volcano plot showing associations between recurrently mutated genes and the NeoAg^hi^CD8^low^ phenotype. The x axis shows log₂ odds ratio, and the y axis shows −log₁₀(P value) from Fisher’s exact tests. Dot size indicates the percentage of samples altered. Red points indicate features significant at FDR < 0.1. **f,** Schematic of recurrent CNA clustering strategy and volcano plot showing associations between recurrent CNA clusters and the NeoAg^hi^CD8^low^ phenotype. The x axis shows log₂ odds ratio, and the y axis shows −log₁₀(*P* value) from Fisher’s exact tests. Dot size indicates the percentage of samples altered. Red points indicate CNA clusters significant at FDR < 0.1. **g,** Relative importance of *TP53* mutation, *ELF3* mutation and CNA Cluster 1 for predicting the NeoAg^hi^CD8^low^ phenotype across three modeling approaches: logistic regression, random forest permutation importance and SHAP analysis. **h,** Analysis of ecDNA in TCGA BLCA whole-genome sequencing data using the AmpliconArchitect pipeline. The stacked bar plot shows the proportion of ecDNA-positive and ecDNA-negative tumors harboring CNA Cluster 1. Odds ratio and *P* value were calculated using a chi-square test. **i,** Association of ecDNA positivity and CNA Cluster 1 with the NeoAg^hi^CD8^low^ phenotype. Odds ratios and *P* values are shown relative to ecDNA-negative tumors and tumors without CNA Cluster 1, respectively.

Consistent with their high neoantigen burden, NeoAg^hi^CD8^low^ tumors also exhibited elevated tumor mutational burden (TMB) (**Supplementary Fig. 1b**). Mutational signature analysis showed that APOBEC3-associated signatures (SBS2/SBS13) were the most strongly enriched signatures in NeoAg^hi^CD8^low^ tumors (**Fig. 1c; Supplementary Fig. 1c, d)**. This was notable because APOBEC3 activity is often linked to immune-inflamed tumor contexts^25^, suggesting that NeoAg^hi^CD8^low^ tumors represent a distinct immune-discordant state within APOBEC3-mutagenized bladder cancer.

To identify transcriptional programs associated with this state, we performed pathway-level analysis of differentially expressed genes. Among the most strongly downregulated programs were pathways involved in tumor–immune engagement (**Supplementary Fig. 1e**). Guided by these findings, we examined a curated immune-engagement gene signature spanning three major components of tumor immune visibility: cytokines and chemokines involved in immune-cell recruitment^26^; antigen-presentation machinery, including HLA class I genes, *B2M*, *TAP1, TAPBP, TAPBPL*, and *PSMB8/9*^27,28^; and epithelial immunoregulatory checkpoint molecules, including *CD274 (PD-L1), CD80, CD86, PDCD1LG2 (PD-L2),* and *CD47*^29–33^. These programs were coordinately suppressed in NeoAg^hi^CD8^low^ tumors (**Fig. 1d**). This phenotype was not explained by somatic HLA loss, and NeoAg^hi^CD8^low^ tumors did not show evidence of inferior predicted neoantigen quality, arguing against poor antigenicity as the primary explanation for reduced immune infiltration (**Supplementary Fig. 1f, g**). Together, these findings define an immune–neoantigen-discordant tumor state characterized by high antigenic potential but coordinated suppression of tumor-cell-intrinsic immune-engagement programs.

We next asked whether recurrent genomic alterations were associated with the NeoAg^hi^CD8^low^ phenotype. Recurrently mutated driver genes present in at least 10% of tumors showed only modest associations: *TP53* and *ELF3* were nominally associated with NeoAg^hi^CD8^low^ status, but neither remained significant after false-discovery-rate correction (FDR < 0.1; **Fig. 1e**). We therefore examined recurrent focal copy-number alterations (CNAs), restricting the analysis to events present in at least 10% of tumors and organizing them into clusters of co-occurring alterations (**Supplementary Fig. 2a, b; Methods**). Three CNA clusters were significantly associated with the NeoAg^hi^CD8^low^ phenotype, with Cluster 1 showing the strongest positive association (**Fig. 1f**). Across statistical models, Cluster 1 outperformed *TP53* and *ELF3* mutation status as a predictor of NeoAg^hi^CD8^low^ tumors (**Fig. 1g**).

Because focal amplifications can arise within complex structural contexts, including extrachromosomal DNA (ecDNA), we next assessed whether ecDNA was associated with immune–neoantigen discordance. Analysis of whole-genome sequencing data from the BLCA cohort (n = 408), using an AmpliconArchitect pipeline^34^, identified ecDNA in 163 tumors. Cluster 1 amplification was significantly enriched in ecDNA-positive tumors (**Fig. 1h**). In multivariable models, both ecDNA status and Cluster 1 amplification showed comparable associations with the NeoAg^hi^CD8^low^ phenotype, indicating that focal amplification and ecDNA are linked genomic features of this immune–neoantigen-discordant state (**Fig. 1i**).

### NECTIN4 is the dominant 1q23.3 predictor of immune–neoantigen discordance

To identify candidate effectors of the NeoAg^hi^CD8^low^ phenotype, we focused on Cluster 1 amplification events. This cluster encompassed a broad chromosome 1q23 region containing 93 genes. Chromosome 1q23 was among the most frequently amplified regions in bladder cancer, with high-level amplification in approximately 17% of tumors and single-copy gain in approximately 35% of tumors by GISTIC analysis (**Supplementary Fig. 2c**), although its functional contribution has remained undefined.

We integrated matched RNA-sequencing and copy-number data to prioritize genes within this interval whose expression was coupled to amplification and associated with immune–neoantigen discordance. Although many 1q23 genes exhibited copy-number-coupled expression, *NECTIN4* showed the strongest association with the NeoAg^hi^CD8^low^ phenotype in both unadjusted and purity-and ploidy-adjusted analyses (**Fig. 2a**). Moreover, when ecDNA status and *NECTIN4* mRNA expression were modeled jointly, *NECTIN4* emerged as the dominant predictor of the NeoAg^hi^CD8^low^ phenotype, outperforming ecDNA status alone (**Fig. 2b**). These analyses nominated NECTIN4 as the principal candidate effector within the 1q23.3 locus.

**Fig. 2.**
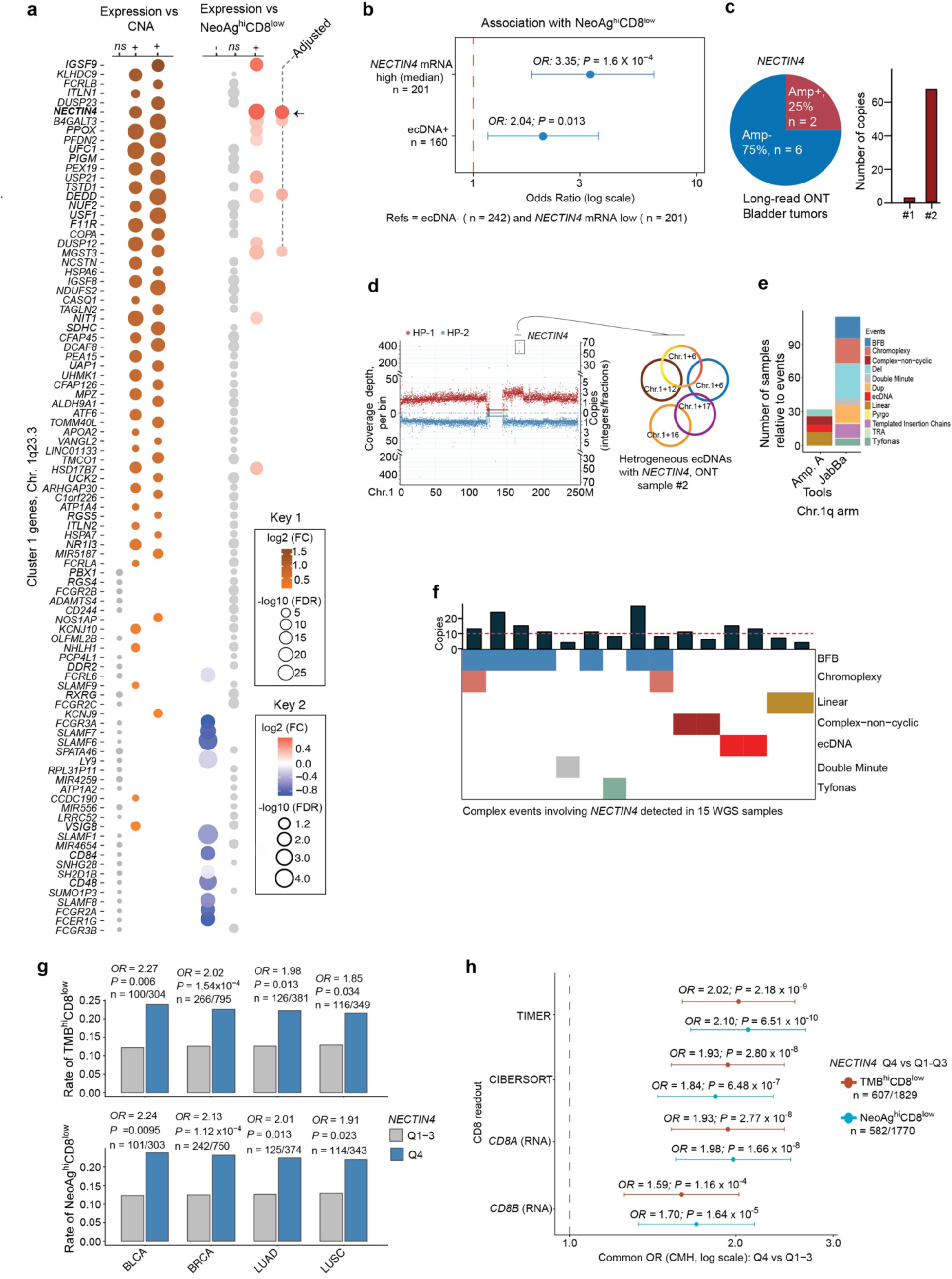
NECTIN4 is the dominant 1q23.3 predictor of the NeoAg^hi^CD8^low^ tumor state. a, Left, associations between CNAs and expression of genes within CNA Cluster 1 at the chromosome 1q23.3 locus. Right, associations between gene expression and the NeoAg^hi^CD8^low^ phenotype. Grey dots denote non-significant associations. The dashed boundary indicates analyses performed using NeoAg^hi^CD8^low^ groups defined after adjustment for tumor purity and ploidy, highlighting *NECTIN4* as the top signal. Dot size represents −log₁₀(FDR), and color indicates the direction and magnitude of association; separate color-scale keys show log₂ fold change. **b,** Multivariable logistic regression models assessing associations with the NeoAg^hi^CD8^low^ phenotype. Forest plots show odds ratios (ORs) for *NECTIN4* mRNA expression and ecDNA status. Points represent ORs, and horizontal lines denote 95% confidence intervals (CIs). The x axis is displayed on a log scale. **c,** Pie chart showing *NECTIN4* amplification in two of eight bladder tumors analyzed by long-read Oxford Nanopore sequencing. The bar plot shows inferred *NECTIN4* copy number in the two amplified samples. **d,** Genome-wide copy-number profile and haplotype-resolved view from a bladder tumor harboring high-level *NECTIN4* amplification. The inset shows a heterogeneous circular amplicon structure involving *NECTIN4*. **e,** Stacked bar plot summarizing the number and classes of complex structural events identified on chromosome 1q using AmpliconArchitect analysis of TCGA bladder cancer whole-genome sequencing data. **f,** Heatmap summarizing complex structural events involving *NECTIN4* identified in whole-genome sequencing samples, including breakage–fusion–bridge, chromothripsis, linear amplification, complex non-cyclic amplification, ecDNA, double-minute and tyfonas events. **g,** Association between high *NECTIN4* expression and TMB^hi^CD8^low^ or NeoAg^hi^CD8^low^ tumor states across bladder (BLCA), breast (BRCA), lung adenocarcinoma (LUAD) and lung squamous cell carcinoma (LUSC) cohorts. High *NECTIN4* expression was defined as quartile 4 (Q4) versus quartiles 1–3 (Q1–3). Odds ratios and *P* values were calculated for each tumor type. **h,** Association between high *NECTIN4* expression and TMB^hi^CD8^low^ or NeoAg^hi^CD8^low^ tumor states across independent CD8⁺ T-cell inference methods, including TIMER, CIBERSORT and RNA-based CD8A and CD8B readouts. Forest plot shows common odds ratios estimated using Cochran–Mantel–Haenszel models across tumor types; horizontal lines denote 95% CIs. The x axis is displayed on a log scale. P values are two-sided, and sample sizes for pooled analyses are indicated.

We next examined whether *NECTIN4* amplification occurred within complex structural contexts. In a subset of eight bladder tumors profiled by long-read Oxford Nanopore sequencing, two harbored *NECTIN4*-region amplification. One tumor showed approximately 68 copies of the *NECTIN4* region together with rearrangements involving multiple chromosomes, consistent with a complex amplified structure compatible with ecDNA or other rearrangement-driven amplification mechanisms (**Fig. 2c-e**). In TCGA whole-genome sequencing data, complex rearrangements were also observed around the *NECTIN4* locus. Using JaBbA^35^, we identified more than ten *NECTIN4*-region copies in ten tumors, further supporting that *NECTIN4* amplification can arise through complex structural mechanisms, including ecDNA-like amplification or breakage–fusion–bridge cycles (**Fig. 2f**).

We further found that *NECTIN4* expression was modulated by promoter methylation, particularly in tumors with copy-number gain, indicating an additional regulatory layer. Across bladder, breast and lung cancers, recurrent *NECTIN4* amplification was accompanied by high median *NECTIN4* mRNA expression (**Supplementary Fig. 3a**). In these malignancies, increased *NECTIN4* copy number was significantly associated with elevated *NECTIN4* expression, whereas promoter methylation further stratified expression among copy-number-gained tumors (**Supplementary Fig. 3b, c**).

Extending the NeoAg^hi^CD8^low^ framework to breast and lung cancers revealed that tumors in the highest quartile of *NECTIN4* expression were significantly enriched for the NeoAg^hi^CD8^low^ phenotype compared with tumors in quartiles 1–3. Similar results were observed for TMB^hi^CD8^low^ tumors (**Fig. 2g**), and these findings were robust across multiple CD8 readouts (**Fig. 2h**). Collectively, these results identify amplification-associated *NECTIN4* overexpression as a recurrent marker of CD8⁺ T-cell paucity in tumors with high neoantigen or mutational burden, motivating functional interrogation of its role in shaping tumor–immune interactions.

### NECTIN4 promotes a T-cell-poor tumor microenvironment and resistance to PD-1 blockade

To determine whether tumor-intrinsic NECTIN4 overexpression can modulate the immune microenvironment in an immunogenic setting, we generated syngeneic MB49 bladder cancer cells constitutively expressing human NECTIN4 and implanted them, alongside vector controls, into immunocompetent C57BL/6 mice (**Fig. 3a, b; Methods**). MB49 tumors are immunologically inflamed, neoantigen-rich and responsive to anti–PD-1 therapy^36–38^, providing a sensitized background in which immune suppression can be readily detected.

**Fig. 3.**
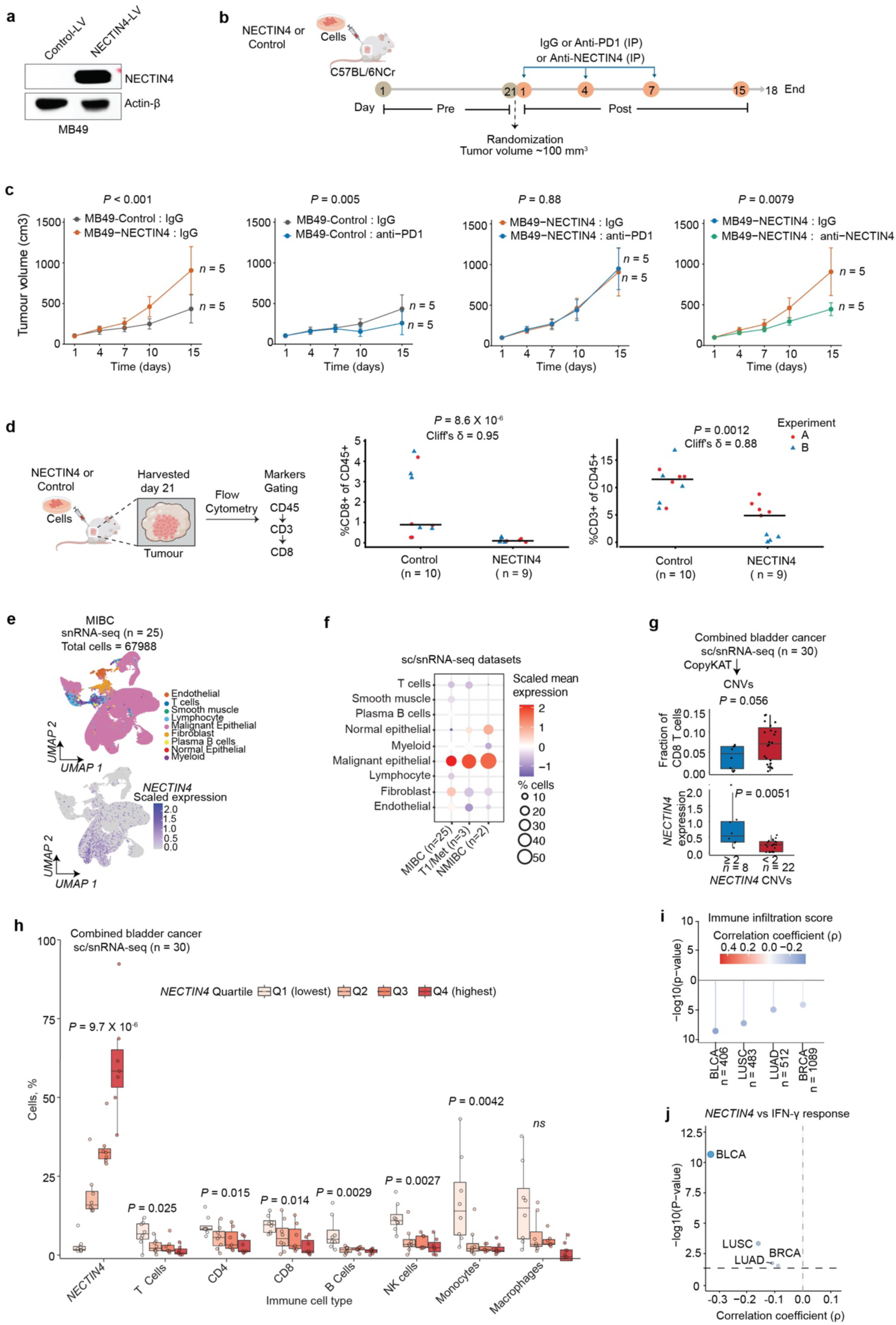
NECTIN4 overexpression induces intratumoral T-cell paucity in vivo and is associated with reduced immune infiltration in human tumors. a, Immunoblot analysis of human NECTIN4 protein in MB49 cells transduced with a human NECTIN4 lentiviral vector or control vector. Actin-β was used as a loading control. **b,** Schematic of syngeneic tumor implantation and antibody-treatment schedule. C57BL/6NCr mice were implanted subcutaneously with parental MB49, control-vector MB49 or NECTIN4-expressing MB49 cells. Tumors were allowed to grow to approximately 100 mm³ before randomization, after which mice received intraperitoneal control IgG, anti–PD-1 or NECTIN4-targeting antibody according to the indicated dosing schedule. **c,** Tumor-growth analysis of parental MB49, control-vector MB49 and NECTIN4-expressing MB49 tumors treated with control IgG, anti–PD-1 or NECTIN4-targeting antibody. Tumor growth was analyzed through day 15, before tumor-burden-related dropout occurred in control groups. Data are shown as mean ± s.e.m.; *P* values indicate planned contrasts of tumor-growth slopes from linear mixed-effects models. **d,** Schematic of syngeneic tumor implantation and flow-cytometric immune profiling. Mice were euthanized three weeks after implantation, and tumors were analyzed using a flow-cytometry panel including CD45, CD3, CD4 and CD8. Right, combined analysis of intratumoral CD3⁺ and CD8⁺ T-cell frequencies across two independent experiments. Each dot represents an individual mouse; horizontal lines indicate medians. *P* values were calculated using two-sided Mann–Whitney U tests. Cliff’s δ values are shown. Flow-cytometry gating and independently reproduced experiment-level results are provided in **Supplementary Data Fig. 1.** **e,** UMAP analysis of bladder cancer single-cell and single-nucleus RNA-sequencing datasets. The upper panel shows annotated major cell types, and the lower panel shows scaled *NECTIN4* expression. **f,** Dot plot showing scaled mean *NECTIN4* expression and the percentage of *NECTIN4*-expressing cells across major cell types in three bladder cancer single-cell or single-nucleus RNA-sequencing datasets. **g,** CopyKAT-inferred *NECTIN4* copy number in combined bladder cancer single-cell and single-nucleus RNA-sequencing datasets. Box plot shows CD8⁺ T-cell abundance in tumors with inferred *NECTIN*4 amplification compared with non-amplified tumors. *P* value was calculated using a two-sided Mann–Whitney U test. **h,** Relative abundance of immune-cell populations stratified by malignant-cell *NECTIN4* expression quartile in combined bladder cancer single-cell and single-nucleus RNA-sequencing datasets. *P* values were calculated using Kruskal–Wallis tests. **i,** Associations between *NECTIN4* expression and immune infiltration scores inferred from bulk RNA-sequencing across cancer types with high *NECTIN4* expression. The y axis shows −log₁₀(FDR-adjusted *P* value), and dot color indicates Spearman’s correlation coefficient. **j,** Associations between *NECTIN4* expression and IFN-γ response scores across cancer types. The y axis shows −log₁₀(FDR-adjusted P value), and the x axis shows Spearman’s correlation coefficient.

Longitudinal mixed-effects analysis of tumor growth through day 15, before tumor-burden-related dropout occurred in control groups, showed that NECTIN4-expressing tumors grew significantly faster than vector-control tumors under IgG treatment (**Fig. 3b, c**). Anti–PD-1 treatment significantly slowed the growth of vector-control tumors but had no significant effect on MB49-NECTIN4 tumors, suggesting that NECTIN4 can blunt the antitumor effect of PD-1 blockade in this model. Conversely, treatment of MB49-NECTIN4 tumors with a NECTIN4-targeting antibody significantly reduced tumor-growth rate relative to IgG-treated MB49-NECTIN4 tumors, further supporting a functional contribution of NECTIN4 to this phenotype (**Fig. 3c**).

We hypothesized that NECTIN4 promotes accelerated tumor growth and PD-1 resistance, at least in part, by imposing a T-cell-poor tumor microenvironment. To test this, we performed immune profiling three weeks after implantation (**Fig. 3d**). NECTIN4-expressing tumors exhibited reduced immune infiltration, with lower fractions of CD3⁺ and CD8⁺ T cells compared with vector-control tumors (**Fig. 3d; Supplementary Data Fig. 1a–c**), as well as reduced CD4⁺ T cells (**Supplementary Data Fig. 1c**). Together, these data demonstrate that enforced NECTIN4 expression is sufficient to promote a T-cell-poor tumor microenvironment and resistance to PD-1 blockade in an otherwise immunogenic model.

To assess whether high *NECTIN4* expression is associated with reduced immune infiltration in human bladder cancer, we analyzed tumor single-cell (scRNA-seq) and single-nucleus (snRNA-seq) datasets. These included newly generated scRNA-seq data from non–muscle-invasive bladder cancer (NMIBC; n = 2), newly generated snRNA-seq data from high-risk NMIBC and one metastatic lesion (n = 3), and a publicly available snRNA-seq dataset of muscle-invasive bladder cancer (MIBC; n = 25)^39^ (**Fig. 3e; Supplementary Fig. 4a**). Across NMIBC and MIBC samples, *NECTIN4* expression was largely confined to malignant epithelial cells (**Fig. 3f**). Using CopyKAT^40^ to infer copy-number alterations in sc/snRNA-seq, we identified tumors harboring *NECTIN4* amplification and observed a reduction in CD8⁺ T-cell abundance in these samples (**Fig. 3g**).

We further stratified all 30 tumors into quartiles according to malignant-cell *NECTIN4* expression. Tumors in the highest *NECTIN4* expression quartile showed significantly reduced infiltration of CD8⁺ and CD4⁺ T cells, as well as reduced abundance of additional immune populations, including monocytes and NK cells (**Fig. 3h**). In independent TCGA bulk RNA-sequencing cohorts, elevated *NECTIN4* expression was consistently associated with reduced immune infiltration and diminished IFN-γ signaling across the four cancer types examined (**Fig. 3i, j**). These inverse relationships were consistent across three independent immune-deconvolution methods (**Supplementary Fig. 4b**). To extend these observations beyond bladder cancer at single-cell resolution, we analyzed an independent breast cancer scRNA-seq cohort^41^ (n = 26), in which *NECTIN4* expression was also negatively associated with multiple anti-tumor immune populations (**Supplementary Fig. 4c, d**). Moreover, analysis of public Visium spatial transcriptomics dataset showed that CD8⁺ T cells were spatially excluded from *NECTIN4*-enriched malignant niches (**Supplementary Fig. 4e–g**).

Given reported TIGIT-mediated interactions among related NECTIN family members^42^, we asked whether the NECTIN4-associated immune-cold state reflected a similar lymphocyte checkpoint interaction. Instead, *NECTIN4*-high tumors showed inverse associations with immune-checkpoint receptor genes, including *TIGIT*; *TIGIT*-expressing T cells were not enriched near NECTIN4-expressing malignant cells; and cell–cell interaction analysis did not support direct NECTIN4–lymphocyte engagement (**Supplementary Fig. 5a–e**).

Together, these data show that NECTIN4 overexpression is sufficient to promote a T-cell-poor, PD-1-resistant tumor microenvironment in vivo, and that human tumor profiling across cancers links high *NECTIN4* expression to reduced cytotoxic T-cell infiltration, spatial T-cell exclusion, and immune-cold tumor states. Consistent with this model, high *NECTIN4* expression was also recurrently associated with poorer outcomes across multiple immune checkpoint blockade–treated cohorts (**Supplementary Fig. 6**).

### NECTIN4 enforces tumor-intrinsic immune cloaking through DDR1–SHP2-mediated STAT1 suppression

We next investigated the molecular mechanisms by which NECTIN4 promotes an immune-cold tumor state. In bladder cancer sc/snRNA-seq datasets and the TCGA cohort, NECTIN4-positive epithelial cells and NECTIN4-high tumors exhibited marked downregulation of antigen presentation, T-cell migration, chemokine responsiveness, granulocyte recruitment, and broader tumor-intrinsic programs that facilitate immune-cell activation and trafficking (**Fig. 4a**; **Supplementary Fig. 7a**). Consistent with the immune-engagement programs associated with the immune–neoantigen-discordant state (**Fig. 1d**), NECTIN4-positive epithelial cells showed coordinated suppression of immune-recruiting cytokines and chemokines, antigen-presentation machinery, and epithelial immunoregulatory checkpoint molecules (**Fig. 4b**). These programs were also inversely associated with *NECTIN4* expression in TCGA bladder cancer and showed similar patterns in breast and lung cancers (**Supplementary Fig. 7b, c**). Together, these findings link NECTIN4 to the coordinated multi-arm suppressive phenotype observed in NeoAg^hi^CD8^low^ tumors (**Fig. 1d**). Because this program occurs in neoantigen-rich tumors and includes suppression of immune-recruiting signals, antigen presentation, and immune-checkpoint ligands, we refer to this state as tumor-intrinsic immune cloaking, distinguishing it from general immune exclusion or immune-deficient states.

**Fig. 4.**
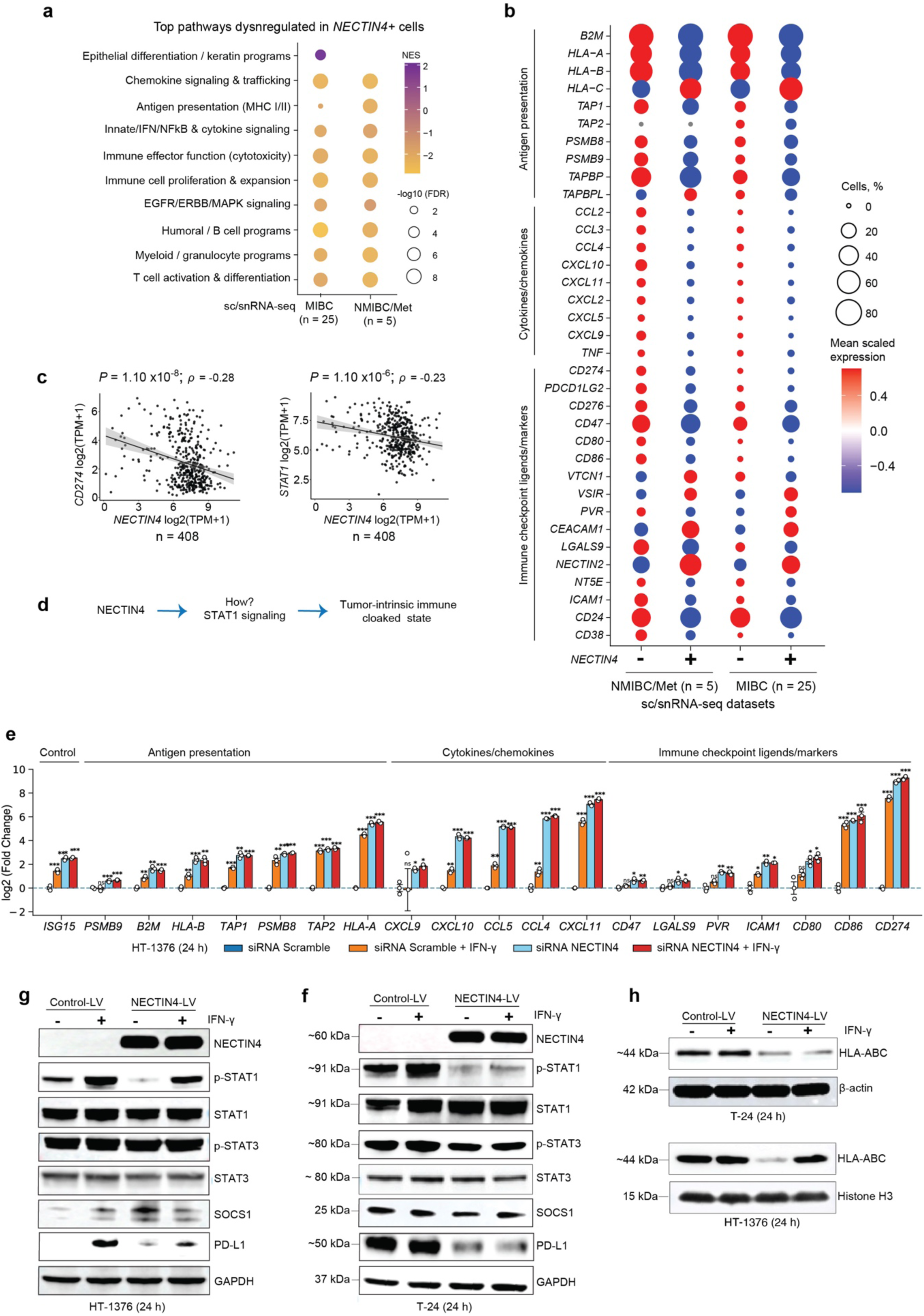
NECTIN4 inhibits STAT1 signaling to attenuate antigen presentation and inflammatory programs. a, Top pathways differentially enriched in *NECTIN4⁺* versus *NECTIN4⁻* epithelial cells across bladder cancer single-cell and single-nucleus RNA-sequencing datasets, including MIBC (n = 25) and NMIBC/metastatic tumors (NMIBC/Met; n = 5). Dot color represents normalized enrichment score (NES), and dot size indicates −log₁₀(FDR). **b,** Dot plot showing expression of antigen-presentation genes, cytokines and chemokines, and immune-checkpoint ligand/marker genes in epithelial cells from bladder cancer single-cell and single-nucleus RNA-sequencing datasets, stratified by *NECTIN4* expression. Dot color denotes mean scaled expression, and dot size represents the percentage of epithelial cells expressing each gene. **c,** Scatter plots showing Spearman rank correlations between *NECTIN4* mRNA expression and *CD274* (PD-L1; left) or *STAT1* (right) in TCGA bladder cancer tumors. Shaded areas indicate 95% confidence intervals. **d,** Schematic model illustrating the hypothesis that *NECTIN4* suppresses *STAT1* signaling to promote a tumor-intrinsic immune-cloaked state. **e,** qPCR analysis of immune-engagement genes in HT-1376 cells following siRNA-mediated NECTIN4 depletion, with or without IFN-γ stimulation. Genes are grouped by antigen presentation, cytokines/chemokines and immune-checkpoint ligands/markers. Data are shown as log₂ fold change relative to control after HPRT normalization. Bars show mean ± s.e.m., and dots indicate biological replicates. Asterisks indicate two-sided Welch’s t-tests versus control: ns, not significant; \**P* < 0.05, \*\**P* < 0.01 and \*\*\**P* < 0.001. **f,** Immunoblot analysis of T24 cells stably transduced with control lentiviral vector or NECTIN4 lentiviral vector, with or without IFN-γ stimulation, assessing NECTIN4 expression, STAT1 and STAT3 phosphorylation, total STAT1 and STAT3, SOCS1 and PD-L1. GAPDH was used as a loading control. **g,** Immunoblot analysis of HT-1376 cells stably transduced with control lentiviral vector or NECTIN4 lentiviral vector, with or without IFN-γ stimulation, assessing NECTIN4 expression, STAT1 and STAT3 phosphorylation, total STAT1 and STAT3, SOCS1 and PD-L1. GAPDH was used as a loading control. **h,** Immunoblot analysis of HLA-ABC expression in T24 and HT-1376 cells stably transduced with control lentiviral vector or NECTIN4 lentiviral vector, with or without IFN-γ stimulation. β-actin and histone H3 were used as loading controls, as indicated.

We therefore asked whether this coordinated suppressive state converges on a common tumor-intrinsic signaling mechanism. STAT1 phosphorylation is a central signaling node controlling transcriptional activation of inflammatory cytokines, antigen-presentation machinery and select immune-checkpoint genes^43^. Consistent with this role, *NECTIN4* levels were negatively correlated with *STAT1* mRNA, which can reflect STAT1-dependent transcriptional feedback, and with the STAT1 target *CD274* (PD-L1) in bladder tumors (**Fig. 4c**). We therefore hypothesized that NECTIN4 suppresses tumor-cell-intrinsic immune signaling through inhibition of STAT1 activity (**Fig. 4d**).

To test this, we performed mechanistic perturbation experiments in HT-1376 cells, which express endogenous NECTIN4 (**Supplementary Fig. 8a**). siRNA-mediated NECTIN4 depletion increased the expression of multiple tumor-cell immune-engagement genes, both at baseline and after IFN-γ stimulation, spanning antigen presentation, cytokine/chemokine signaling and immune-checkpoint/adhesion programs (**Fig. 4e**). These findings indicate that endogenous NECTIN4 restrains immune-visibility programs at the transcript level, consistent with its role in promoting tumor-intrinsic immune cloaking.

We next asked whether this transcriptional effect was reflected at the protein and signaling levels. In HT-1376 cells, NECTIN4 knockdown increased STAT1 phosphorylation and PD-L1 protein expression, whereas NECTIN4 overexpression attenuated IFN-γ–induced STAT1 phosphorylation and reduced PD-L1 expression, a tractable downstream readout of STAT1 transcriptional activity^43^ (**Fig. 4g; Supplementary Fig. 8b**). To determine whether this effect was reproducible in a complementary NECTIN4-negative model, we stably expressed NECTIN4 in T24 cells. Consistent with the HT-1376 findings, NECTIN4 expression in T24 cells diminished IFN-γ–induced STAT1 activation and reduced downstream immune-engagement proteins, including PD-L1 (**Fig. 4f**). HLA-ABC was also reduced in NECTIN4-overexpression models and showed moderate upregulation in the HT-1376 NECTIN4-knockdown model (**Fig. 4h; Supplementary Fig. 8b**). In contrast, STAT3 phosphorylation and SOCS1 levels were not consistently altered across conditions, suggesting pathway selectivity toward STAT1 rather than broad suppression of JAK–STAT signaling. Notably, NECTIN4 ectopic expression did not affect cell proliferation in T24 cells (**Supplementary Fig. 8c**). Altogether, these results indicate that NECTIN4 mediates intrinsic immune suppression through selective inhibition of STAT1 signaling.

To elucidate how NECTIN4 transduces signals from the plasma membrane to STAT1, we performed affinity purification of Myc-and DDK-tagged NECTIN4 from HT-1376 cells followed by mass spectrometry (**Fig. 5a**). This approach identified 30 proteins consistently recovered in both pulldowns, including multiple membrane-associated factors (**Fig. 5b, c**). Among these was discoidin domain receptor 1 (DDR1), a receptor tyrosine kinase reported to promote STAT1 dephosphorylation through SHP2 phosphatase activity^44^ and has been implicated in immune suppression^45^. The NECTIN4–DDR1 interaction was further confirmed by co-immunoprecipitation (**Fig. 5d**). Consistent with these findings, analysis of bladder cancer sc/snRNA-seq datasets revealed that *DDR1* expression was significantly reduced in NECTIN4-low epithelial cells, whereas *DDR2*—reported to counteract DDR1 activity^45^—was enriched in NECTIN4-low relative to NECTIN4-high cells (**Fig. 5e**).

**Fig. 5.**
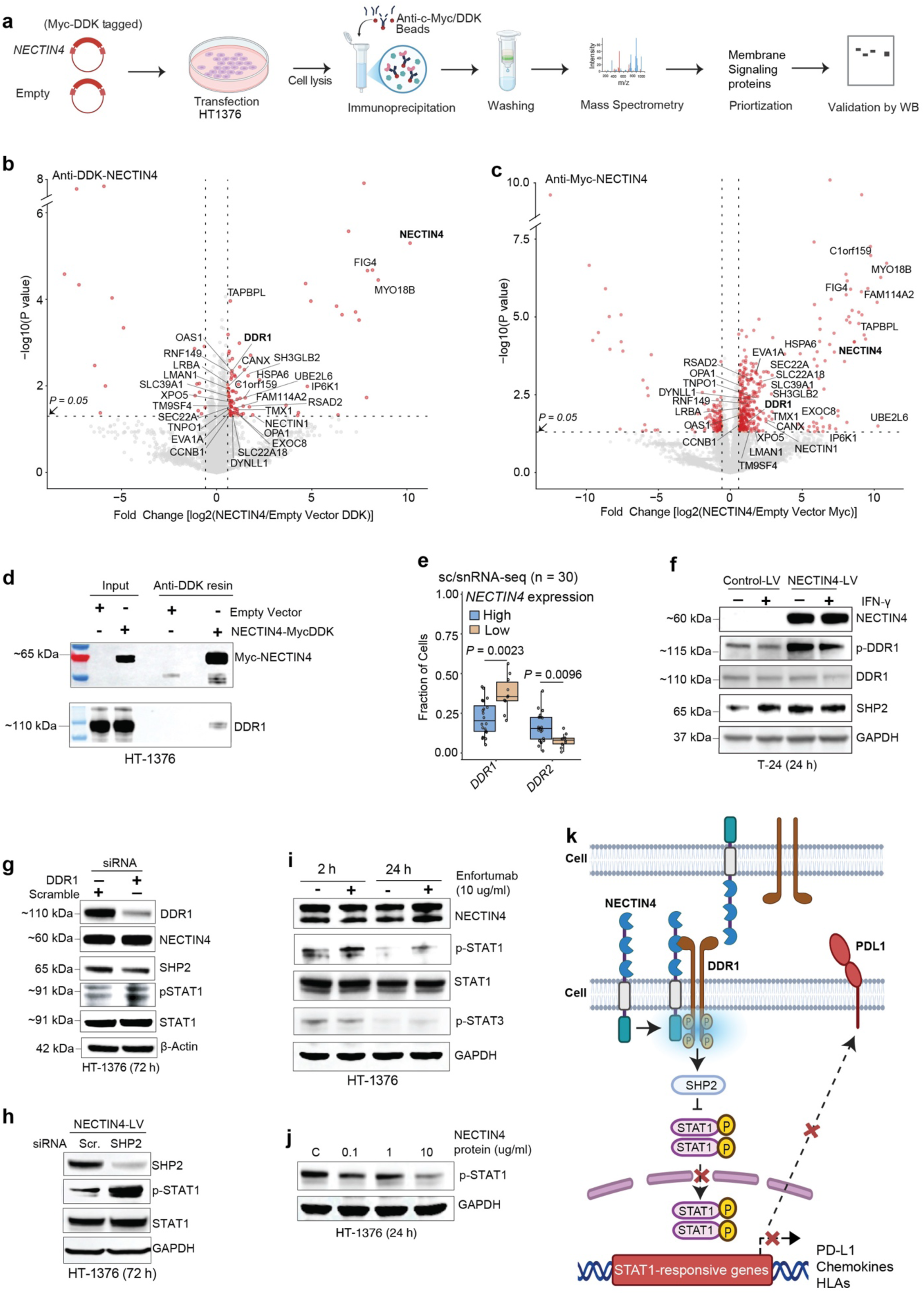
DDR1 couples NECTIN4 to STAT1 inhibition. a, Schematic of affinity purification–mass spectrometry workflow. HT-1376 cells were transfected with Myc-DDK-tagged NECTIN4 or empty vector, followed by immunoprecipitation using anti-Myc or anti-DDK beads, mass spectrometry analysis, prioritization of membrane-associated signaling proteins and validation by immunoblotting. **b,c,** Volcano plots showing proteins enriched in anti-DDK–NECTIN4 (**b**) and anti-Myc–NECTIN4 (**c**) pulldowns compared with empty-vector controls. The x axis shows log₂ fold change, and the y axis shows −log₁₀(P value). Proteins significantly enriched in both pulldowns are highlighted, with selected membrane-associated proteins labeled, including NECTIN4 and DDR1. **d,** Co-immunoprecipitation followed by immunoblotting confirming the interaction between Myc-DDK-tagged NECTIN4 and endogenous DDR1 in HT-1376 cells. **e,** Fraction of malignant epithelial cells expressing DDR1 or DDR2 in *NECTIN4*-high and *NECTIN4*-low tumors across bladder cancer single-cell and single-nucleus RNA-sequencing datasets. Tumors were stratified by *NECTIN4* expression. *P* values were calculated using two-sided Wilcoxon rank-sum tests. **f,** Immunoblot analysis of T24 cells stably transduced with control lentiviral vector or NECTIN4 lentiviral vector, with or without IFN-γ stimulation, assessing NECTIN4, phosphorylated DDR1, total DDR1 and SHP2. GAPDH was used as a loading control. **g,** Immunoblot analysis of HT-1376 cells following siRNA-mediated DDR1 knockdown, assessing DDR1, NECTIN4, SHP2, phosphorylated STAT1 and total STAT1. β-actin was used as a loading control. **h,** Immunoblot analysis of HT-1376 NECTIN4-overexpressing cells following siRNA-mediated SHP2 knockdown, assessing SHP2, phosphorylated STAT1 and total STAT1. GAPDH was used as a loading control. **i,** Immunoblot analysis of HT-1376 cells treated with enfortumab for 2 h or 24 h, assessing NECTIN4, phosphorylated STAT1, total STAT1 and phosphorylated STAT3. GAPDH was used as a loading control. **j,** Immunoblot analysis of phosphorylated STAT1 in HT-1376 cells treated with recombinant NECTIN4 protein at the indicated concentrations for 24 h. GAPDH was used as a loading control. **k,** Proposed model in which extracellular NECTIN4 engagement activates DDR1 at the cell surface, promoting SHP2-dependent inhibition of STAT1 signaling and thereby reducing STAT1-responsive immune-engagement programs, including antigen presentation, chemokines and PD-L1.

We next evaluated the effect of NECTIN4 on DDR1 activity. NECTIN4 overexpression increased DDR1 phosphorylation and elevated levels of the phosphatase SHP2 (**Fig. 5f**; **Supplementary Fig. 8d**). In HT-1376 cells with endogenous NECTIN4 expression, *DDR1* knockdown increased STAT1 phosphorylation (**Fig. 5g**). Consistently, both *DDR1* and SHP2 knockdown increased STAT1 phosphorylation in NECTIN4-overexpression models (**Fig. 5h; Supplementary Fig. 8e, f**). These results are consistent with DDR1 recruiting SHP2 to dephosphorylate STAT1^44,46^, linking NECTIN4–DDR1 engagement to STAT1 suppression.

We therefore asked whether STAT1 suppression reflects extracellular NECTIN4 engagement. Treatment with a NECTIN4-blocking antibody was sufficient to increase STAT1 phosphorylation (**Fig. 5i**), whereas recombinant NECTIN4 reduced STAT1 phosphorylation (**Fig. 5j**). These findings indicate that extracellular NECTIN4 engagement suppresses STAT1 signaling. This conclusion is consistent with enrichment of GFP-tagged NECTIN4 at cell–cell boundaries (**Supplementary Fig. 8g**) and with the predominance of epithelial–epithelial interactions observed in single-cell interaction analyses (**Supplementary Fig. 5d**). Overall, these data support a model in which extracellular NECTIN4 engagement activates DDR1 at the cell surface, promoting SHP2-dependent inhibition of STAT1 signaling and thereby enforcing tumor-intrinsic immune cloaking (**Fig. 5k**).

### APOBEC3 mutagenesis is linked to recurrent 1q23.3 amplification

Having established amplification-driven *NECTIN4* overexpression as a recurrent feature of immune–neoantigen discordance, we next asked whether specific mutational processes were associated with the repeated emergence of 1q23.3 amplification. Using machine-learning approaches (**Methods**), we assessed associations between recurrently amplified genes and common mutational signatures present in at least 10% of samples. Across models, four SBS signatures — SBS1, SBS2, SBS5, and SBS13— emerged as the strongest predictors of recurrent amplifications (**Supplementary Data Tables 1 and 2**). Univariate analyses revealed that distinct CNAs were preferentially associated with specific mutational processes (**Fig. 6a**). Notably, amplification of chromosome 1q regions, including the *NECTIN4-*containing 1q23.3 locus, was strongly associated with SBS2 and SBS13, whereas amplifications on chromosomes 3 and 6 were primarily linked to SBS1 (**Fig. 6a, b**). Because SBS2 and SBS13 are canonical APOBEC3-associated signatures and represent dominant sources of somatic single-nucleotide variation in hypermutated bladder, lung, and breast cancers, these findings implicated APOBEC3 mutagenesis as a genomic correlate of recurrent *NECTIN4* amplification^14–17^. Consistent with this interpretation, SBS2/SBS13 mutational burden was higher in *NECTIN4*-amplified tumors (**Fig. 6c**), similar to its association with the NeoAg^hi^CD8^low^ phenotype (**Fig. 1c**).

**Fig. 6.**
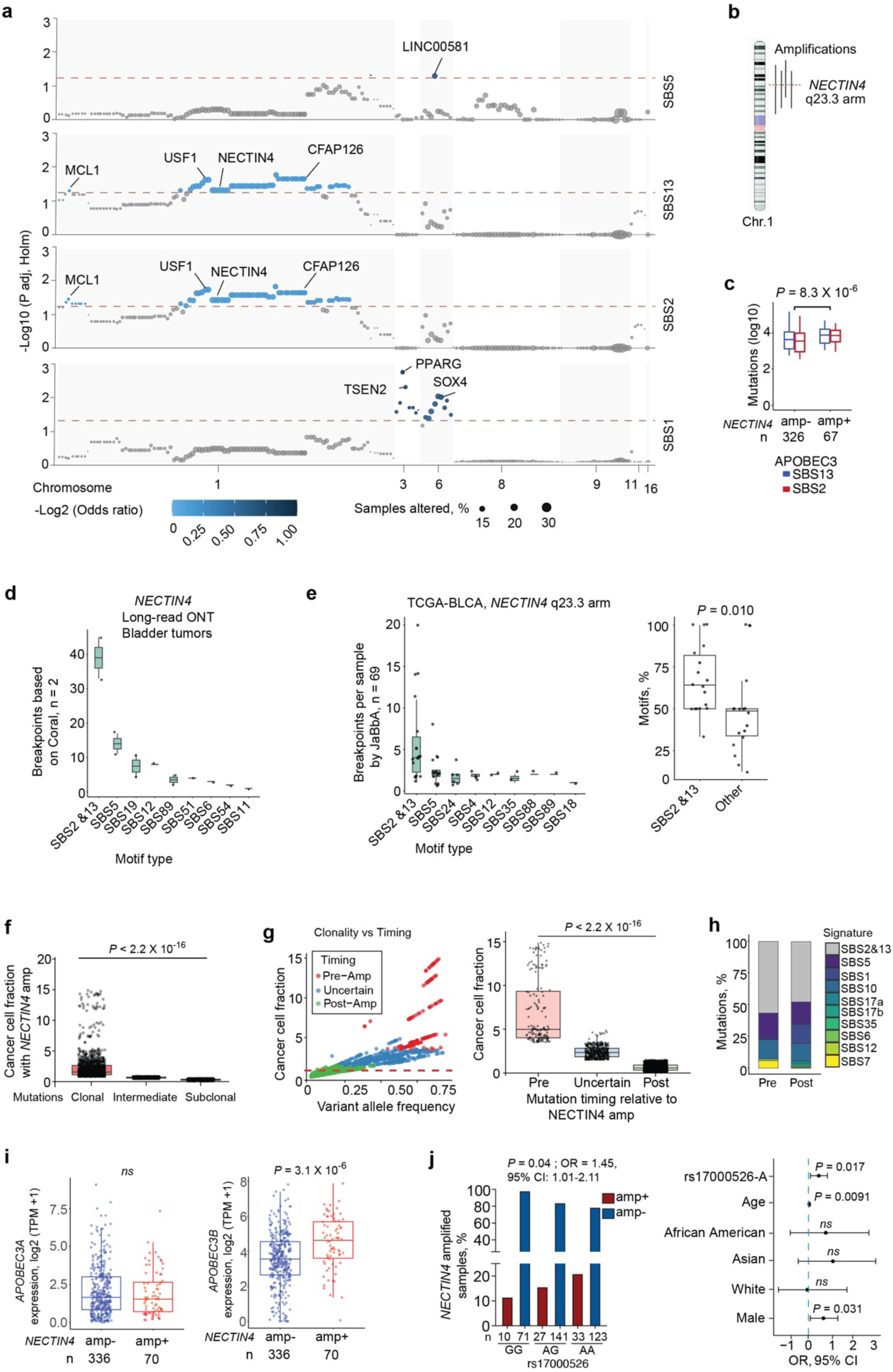
Recurrent amplification of chromosome 1q23.3 harboring *NECTIN4* is associated with APOBEC3-mediated mutagenesis. a, Manhattan plots showing associations between recurrent focal copy-number alterations (CNAs) and mutational signatures prioritized by machine-learning approaches. Associations were tested using univariate logistic regression with Holm correction. The y axis shows −log₁₀(P value), and the x axis indicates genomic position across chromosomes. Dot size represents the frequency of deep amplification or deletion events inferred by GISTIC, and color denotes the log₂ odds ratio. *NECTIN4* amplification and co-amplified genes on chromosome 1q23.3 show strong associations with APOBEC3-associated signatures SBS2 and SBS13. **b,** Chromosome 1 ideogram highlighting the recurrent 1q23.3 amplicon containing *NECTIN4*. **c,** Box plots showing APOBEC3-associated SBS2 and SBS13 mutational burden in *NECTIN4*-amplified and non-amplified tumors. *P* values were calculated using two-sided Mann–Whitney U tests. Boxes indicate medians and interquartile ranges. **d,** Breakpoint-proximal motif distribution in long-read Oxford Nanopore–sequenced bladder tumors. Motifs corresponding to APOBEC3-associated signatures, including SBS2 and SBS13, are shown among breakpoint-associated sequences. **e,** Breakpoint-proximal motif distribution in TCGA bladder cancer whole-genome sequencing data for tumors harboring *NECTIN4* 1q23.3 amplification. Left, motif distribution across mutational signatures. Right, comparison of SBS2/SBS13-associated motif coverage versus other motifs. *P* value was calculated using a two-sided Mann–Whitney U test. **f,** Clonality analysis of somatic mutations located within *NECTIN4* amplicons. Mutations were classified as clonal, intermediate or subclonal based on cancer cell fraction estimates. P value was calculated using a two-sided Kruskal–Wallis test. **g,** Timing analysis of mutations within *NECTIN4* amplicons relative to amplification. Left, variant allele frequency plot used to infer mutation timing categories based on cancer cell fraction and copy-number context. Right, box plot showing cancer cell fraction across pre-amplification, uncertain and post-amplification mutation categories. P value was calculated using a two-sided Kruskal–Wallis test. **h,** Mutational signature composition of somatic mutations within *NECTIN4* amplicons. Stacked bar plots show the relative contribution of mutational signatures among pre-and post-amplification mutations, highlighting APOBEC3-associated SBS2 and SBS13. **i,** Expression of APOBEC3A and APOBEC3B in *NECTIN4*-amplified and non-amplified TCGA bladder tumors. *P* values were calculated using two-sided Mann–Whitney U tests. Boxes indicate medians and interquartile ranges. **j,** Association between germline genetic variation and *NECTIN4* amplification status. Left, percentage of *NECTIN4*-amplified tumors stratified by genotype at rs17000526. Right, forest plot showing odds ratios from a multivariable model including rs17000526 genotype, age, race and sex.

We next asked whether APOBEC3-associated mutational processes were linked to the structural breakpoints underlying *NECTIN4* amplification. Because breakpoint-proximal regions contain limited numbers of substitutions, classical mutational signature deconvolution is underpowered in this setting. We therefore used breakpoint-proximal motif analysis to assess APOBEC3 involvement. APOBEC3-compatible motifs were frequent across recurrent copy-number–altered regions and at chromosome 1 amplification sites, including *NECTIN4*-specific breakpoints identified in long-read Oxford Nanopore–sequenced *NECTIN4*-amplified tumors (**Fig. 6d; Supplementary Fig. 9a**). Similarly, in TCGA bladder cancer whole-genome sequencing data, breakpoint motifs mapped predominantly to APOBEC3-associated signatures genome-wide and at chromosome 1 amplification loci encompassing *NECTIN4* rearrangements (**Fig. 6e; Supplementary Fig. 9b**). *NECTIN4*-containing 1q23.3 locus in breast and lung tumors showed a similar predominance of APOBEC3-associated motif coverage (**Supplementary Fig. 10**).

To examine the evolutionary timing of *NECTIN4* amplification, we analyzed the clonality and timing of somatic mutations within *NECTIN4* amplicons. Mutations mapping to the amplified regions were predominantly clonal relative to the amplification event, indicating that many arose early during tumor evolution (**Fig. 6f**). Variant allele frequency–based timing analysis further showed that a substantial fraction of mutations occurred prior to or during amplification, rather than exclusively after copy-number gain (**Fig. 6g**). Consistent with these observations, mutational signature analysis revealed a strong predominance of APOBEC3-associated signatures (SBS2 and SBS13) among both pre-and post-amplification mutations within *NECTIN4* amplicons (**Fig. 6h**). Together, these results support a model in which APOBEC3-mediated mutagenesis creates a genomic context that facilitates the recurrent emergence of *NECTIN4* amplification during tumor evolution.

APOBEC3 cytidine deaminases have previously been implicated in complex rearrangements and copy number alterations^47^. Supporting this link, *APOBEC3B* expression was positively associated with *NECTIN4* amplification (**Fig. 6i**). Moreover, a germline variant identified by genome-wide association studies and known to influence *APOBEC3B* expression in bladder cancer^48^, was significantly associated with *NECTIN4* amplification (**Fig. 6j**). Because this variant reflects inherited predisposition to altered APOBEC3 activity, these findings provide orthogonal genetic support for an APOBEC3B-linked contribution to recurrent *NECTIN4* amplification. Overall, these data support a model in which APOBEC3-mediated mutagenesis is linked to recurrent 1q23.3 amplification and associated rearrangements, enabling the emergence of NECTIN4-driven tumor-intrinsic immune cloaking in neoantigen-rich tumors.

## Discussion

Our findings reveal a mechanism by which hypermutated tumors mitigate the immunologic liability of high neoantigen burden during tumor evolution. Rather than simply increasing antigenicity, APOBEC3-mediated mutagenesis is linked to recurrent 1q23.3 amplification and N*ECTIN4* overexpression, establishing a tumor-intrinsic program that suppresses immune visibility despite abundant neoantigens. Through inhibition of STAT1 signaling, NECTIN4 attenuates chemokine-mediated immune recruitment, antigen presentation and interferon-inducible immune-engagement programs. These findings connect neoantigen-generating mutational processes, structural genome evolution and tumor-cell-intrinsic immune suppression, providing a mechanistic basis for how neoantigen-rich tumors can remain immunologically silent.

We conceptualize this phenomenon as tumor-intrinsic immune cloaking: a tumor-cell state in which antigenically rich tumors remain immunologically cold by suppressing the programs required for immune engagement. This state is distinct from canonical checkpoint-mediated immune evasion. Whereas checkpoint-dominant tumors are typically inflamed and depend on ongoing immune engagement, tumor-intrinsic immune cloaking prevents immune activation at its source. This distinction provides a potential explanation for the NECTIN4-associated resistance to PD-1 blockade observed in our model: the immune-engagement programs required for productive antitumor immunity are suppressed before checkpoint inhibition can be effective. Consistent with this model, NECTIN4 suppresses not only antigen presentation and T-cell recruitment programs but also PD-L1 expression, distinguishing cloaking from adaptive feedback inhibition in inflamed tumors.

Importantly, this cloaked state appears reversible. In tumor-cell models, NECTIN4 blockade restored STAT1 phosphorylation, suggesting that NECTIN4-dependent suppression of immune signaling can be relieved. Because STAT1 regulates antigen-presentation machinery, inflammatory cytokines and PD-L1 expression, restoration of STAT1 activity may shift immune-cold, neoantigen-rich tumors toward a more immune-engaged and checkpoint-regulated state. In vivo, NECTIN4 overexpression imposed T-cell paucity and resistance to PD-1 blockade, whereas NECTIN4-targeting antibody treatment reduced tumor growth, supporting the functional relevance of this pathway in an immunocompetent setting.

Clinically, NECTIN4 is already targetable, most notably through enfortumab vedotin in urothelial carcinoma. Although enfortumab vedotin has cytotoxic activity through its antibody–drug conjugate payload, our findings suggest that NECTIN4 biology may also intersect with tumor-intrinsic immune signaling. This model provides a potential biological rationale for the clinical benefit observed with enfortumab vedotin combined with PD-1/PD-L1 blockade in metastatic urothelial carcinoma^49^, including its superiority over checkpoint blockade alone in clinical trials^49–51^. Future studies should determine whether NECTIN4-targeting strategies can reverse tumor-intrinsic immune cloaking and improve checkpoint responsiveness.

Across bladder, breast and lung cancers, elevated *NECTIN4* expression was consistently associated with reduced immune infiltration and immune–neoantigen discordance. In these tumors, *NECTIN4* upregulation arises predominantly through recurrent 1q23.3 copy-number gain, with additional modulation by promoter methylation. These findings suggest that mutational processes not only generate neoantigens but can also shape structural genome evolution toward immune-evasive states. In this model, APOBEC3-associated mutagenesis and recurrent 1q23.3 amplification converge to enable NECTIN4-driven immune cloaking, linking neoantigen-generating mutational processes to copy-number-mediated immune escape. Notably, *NECTIN4* is physiologically expressed in the placenta (**Supplementary Data Fig. 2**), a tissue characterized by immune tolerance during gestation^52^, raising the possibility that tumors co-opt developmental immune-regulatory programs to achieve immune invisibility.

At the mechanistic level, NECTIN4 engages DDR1, a receptor tyrosine kinase previously implicated in immune exclusion^45,53^. Our findings place DDR1 upstream of SHP2-dependent STAT1 suppression, providing a signaling mechanism by which a cell-surface adhesion molecule can coordinately repress antigen presentation, immune-recruiting cytokines and checkpoint-associated immune-engagement programs. Future work will be required to define how this axis integrates with DDR1-mediated matrix remodeling, stress responses and inflammatory signaling and whether these interactions differ across tumor types.

Our syngeneic NECTIN4 overexpression model provides direct in vivo evidence that NECTIN4 is sufficient to induce intratumoral T cell paucity; however, it does not fully capture the genomic and regulatory complexity of the 1q23.3 amplicons. Future studies will be required to systematically dissect the functional landscape of this locus and determine whether additional co-amplified genes cooperate with NECTIN4 to reinforce tumor-intrinsic immune cloaking. Moreover, studies across additional immunocompetent tumor models will help define the broader role of NECTIN4 in shaping tumor–immune interactions and checkpoint responsiveness.

More broadly, these results illustrate how hypermutated tumors can maintain an immune-cold state despite high neoantigen burden and provide a framework for identifying genome-encoded vulnerabilities in immunologically refractory cancers.

## Materials and Methods

### Ethical statement

Patient recruitment and tissue procurement were conducted under National Cancer Institute, National Institutes of Health (NIH) Institutional Review Board protocols NCT02379429 and NCT04923178. Recombinant DNA experiments were reviewed and approved by the NIH Institutional Biosafety Committee (IBC; RD-24-II-28). Animal studies were performed under NIH Institutional Animal Care and Use Committee (IACUC) protocols GMB-012 and GMB-014, in accordance with the NIH Guide for the Care and Use of Laboratory Animals.

### Human tumor omics datasets and data sources

Both newly generated and publicly available human datasets were analyzed. Newly generated datasets were derived from histologically confirmed urothelial carcinoma samples collected from patients enrolled under National Cancer Institute, National Institutes of Health Institutional Review Board–approved clinical protocols, with written informed consent obtained from all participants. The newly generated cohort comprised 13 bladder tumors from 10 patients, including 8 male and 2 female participants, consistent with the underlying epidemiology of bladder cancer. Basic clinical covariates, including age at diagnosis, stage, primary versus metastatic status, and tumor subtype, are provided in the associated metadata file. Five matched peripheral blood mononuclear cell (PBMC) samples and five matched normal urothelium samples were also collected; normal samples were used for copy-number profiling and structural variant analyses. We generated single-cell and single-nucleus RNA-sequencing data from five bladder tumors (10x Genomics) and Oxford Nanopore long-read sequencing data from eight bladder tumors with five matched PBMC samples (**Supplementary Data Table 3**).

Whole-genome sequencing (WGS) data for 408 TCGA bladder tumors with matched normal blood samples were accessed through dbGaP (project 35035); only a subset of these samples has been reported previously. WGS data were also analyzed for a subset of breast and lung tumors with GISTIC2 calls of 2, indicating NECTIN4 amplification. Additional datasets included TCGA pan-cancer RNA-sequencing and copy-number alteration data (∼11,000 samples), accessed through the UCSC Xena Browser or cBioPortal. We also analyzed single-nucleus RNA-sequencing datasets from 25 bladder tumors (GEO: GSE169379) and 26 breast tumors (GEO: GSE176078), as well as immunotherapy-response datasets from 32 clinical trials (3,579 patients) accessed through the Cancer Immunology Data Engine (CIDE) before October 30, 2025. Detailed experimental and computational procedures are described below. Other genomics datasets used in this study are referenced and described in the relevant methods sections.

### AmpliconSuite-pipeline for ecDNA characterization

To detect complex structural rearrangements, we analyzed TCGA bladder cancer WGS data (n = 408). Tumor BAM files were used as input and processed via the Singularity wrapper with a pre-downloaded AA_DATA_REPO reference directory. Focal amplification structure and ecDNA status were evaluated using AmpliconSuite-pipeline,^34^ which wraps AmpliconArchitect for reconstruction and AmpliconClassifier for mechanistic classification. We processed tumor BAMs through AmpliconSuite (which invokes AmpliconArchitect and AmpliconClassifier) to reconstruct focal amplicon structures and classify circular amplicons consistent with ecDNA. Copy-number inputs were generated with CNVkit^54^ prior to amplicon reconstruction. AmpliconClassifier was then applied to AmpliconArchitect outputs to classify focal amplifications and report ecDNA/BFB-related calls and related annotations.

### Residual-based identification of neoantigen-high, CD8-low tumors

To identify tumors with disproportionately low cytotoxic T-cell infiltration relative to neoantigen burden, we implemented a residual-based regression framework. Matched TCGA bladder cancer samples (n = 406) with available neoantigen load and CD8⁺ T-cell infiltration estimates were analyzed. Precomputed neoantigen loads^17^, derived using affinity-based prediction of mutant 9–10 mer peptides binding to patient-specific HLA alleles with NetMHCpan, were log₂-transformed, and CD8⁺ T-cell abundance was derived from immune deconvolution of Bulk-RNA-seq data using CIBERSORTx^55^. Only samples with complete data were included. The relationship between neoantigen load and CD8⁺ T-cell infiltration was modeled using generalized additive models (GAMs) to account for non-linear effects.

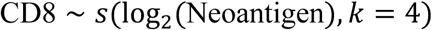

An adjusted model additionally included tumor purity and ploidy as linear covariates. Models were fit using restricted maximum likelihood (REML) as implemented in the *mgcv* R package. Predicted CD8⁺ T-cell values were obtained for each sample, and residuals were calculated as the difference between observed and predicted CD8⁺ T-cell abundance. Tumors were classified as neoantigen-high if their neoantigen burden was greater than or equal to the cohort median. Within this subset, tumors with residual CD8⁺ T-cell values in the lowest quartile (≤25th percentile) were designated as “NeoAg^hi^CD8^low^” tumors, reflecting disproportionately low immune infiltration given high intrinsic immunogenicity. Classification was performed separately for unadjusted and adjusted models to ensure robustness.

### Neoantigen quality assessment

Neoantigen quality was assessed in TCGA bladder cancer samples using the TCGA PanCancer Atlas peptide–MHC binding dataset, generated with NetMHCpan from nonsynonymous somatic mutations identified by MC3 variant calling^4,56^. Predicted mutant peptides of 8–11 amino acids were annotated for HLA class I binding affinity and matched tumor gene expression. Expressed candidate neoantigens were defined as mutant peptide–HLA pairs with predicted binding affinity IC50 <500 nM and source-gene expression>1 TPM, consistent with established neoantigen prediction frameworks. For each sample, we calculated both the number of expressed candidate neoantigens and the fraction of all predicted peptide–HLA pairs meeting these expression and binding criteria. These metrics were used to compare predicted neoantigen quality between NeoAg^hi^CD8^low^ tumors and other tumors within the TCGA bladder cancer cohort.

### Assessment of HLA class I loss and predicted antigenicity

Somatic HLA class I loss was assessed using LOHHLA-derived allele-specific loss calls, HLA homozygosity status and nonsense mutations affecting *HLA-A*, *HLA-B*, *HLA-C* and *B2M*, obtained from a previously described TCGA pan-cancer dataset^57^. Associations with the NeoAg^hi^CD8^low^ phenotype were tested using Fisher’s exact tests for individual loss features and logistic regression models adjusted for tumor purity and ploidy. Composite features included any HLA class I loss, any LOHHLA loss and any nonsense mutation affecting HLA class I genes or *B2M*. P values were corrected for multiple testing using the Benjamini–Hochberg method.

### Association of recurrent genomic alterations and gene expression with the NeoAg^hi^CD8^low^ phenotype

Associations between genomic alterations and the NeoAg^hi^CD8^low^ subset were evaluated using matched somatic mutation, copy-number alteration (CNA), gene expression, and immune data from TCGA bladder cancer samples. For most analyses, we present the unadjusted NeoAg^hi^CD8^low^ subset unless otherwise stated, as it involved fewer sample exclusions; adjusted analyses yielded similar results.

### Somatic mutation analysis

Somatic mutations from TCGA BLCA exomes were obtained from cBioPortal and verified with previous studies^17^. To reduce multiple-testing burden and limit noise from low and rare events, analyses were restricted to genes mutated in ≥10% of samples. Associations between recurrently mutated genes and the NeoAg^hi^CD8^low^ phenotype were assessed using two-sided Fisher’s exact tests. Effect sizes are reported as odds ratios (OR). Multiple testing correction was performed using the Benjamini–Hochberg false discovery rate (FDR) procedure, with FDR < 0.1 considered significant.

### Copy-number alteration clustering and association testing

Genome-wide CNA calls for BLCA were obtained from GISTIC2 analysis of TCGA data. Given the large number of CNA events (>20,000), analyses were restricted to recurrent CNAs present in ≥10% of samples. Recurrent CNAs were grouped by pairwise co-occurrence (Jaccard similarity ≥0.75) and hierarchically clustered, yielding ten CNA clusters representing coordinated genomic alterations. For each cluster, a binary per-sample indicator denoted the presence of any CNA within the cluster. Associations between CNA clusters and the NeoAg^hi^CD8^low^ phenotype were assessed using two-sided Fisher’s exact tests. Effect sizes are reported as odds ratios (OR). Multiple testing correction was performed using the Benjamini–Hochberg false discovery rate (FDR) procedure, with FDR < 0.1 considered significant.

### Multimodel prioritization of CNA Cluster 1

Cluster 1 was prioritized based on its strong association with the NeoAg^hi^CD8^low^ phenotype. Predictive strength relative to TP53 and ELF3 mutation status was evaluated using multivariable logistic regression, random forest (RF) classifiers, and SHAP-based feature attribution^58,59^. Logistic regression models included CNA Cluster 1, TP53 mutation status, and ELF3 mutation status as covariates. Effect sizes are reported as log₂ odds ratios with 95% confidence intervals. RF feature importance was estimated using impurity-based metrics, and SHAP values were used to quantify feature contributions.

### Gene expression analyses within CNA clusters

Gene expression levels for genes within each cluster were evaluated using matched bulk RNA-seq data. Associations between copy number status (GISTIC2-defined amplification versus non-amplification) and gene expression within CNA Cluster 1 were evaluated using two-sided Mann–Whitney U tests. Associations between gene expression and NeoAg^hi^CD8^low^ status were assessed by comparing continuous expression levels between groups using two-sided Mann–Whitney U tests, with false discovery rate correction applied across genes.

### ecDNA analyses with CNA and expression models

Tumors were classified as ecDNA-positive or ecDNA-negative based on the presence of circular amplicons as determined by AmpliconArchitect.^34^ Enrichment of CNA clusters in ecDNA-positive tumors was assessed using chi-square tests. To compare the relative contributions of genomic architecture and gene expression, multivariable logistic regression models were constructed including ecDNA status with CNA Cluster 1 or *NECTIN4* mRNA expression as predictors of the NeoAg^hi^CD8^low^ phenotype. Model coefficients are reported as log₂ odds ratios with 95% confidence intervals.

### Cross-tumor validation of NECTIN4 and TMB^hi^CD8^low^ and NeoAg^hi^CD8^low^ associations

To evaluate generalizability across tumor types, matched neoantigen burden, tumor mutation burden (non-silent mutations) previously computed^4^, gene expression, and CD8⁺ T-cell infiltration estimates were obtained for bladder cancer (BLCA), breast cancer (BRCA), lung adenocarcinoma (LUAD), and lung squamous cell carcinoma (LUSC). These Neoantigen estimates are based on somatic mutations and patient-specific HLA binding affinity^4^. CD8⁺ T-cell abundance was estimated from bulk RNA-seq data using TIMER^60^. TIMER provides tumor purity adjusted estimates. Tumors were stratified by *NECTIN4* expression quartiles within each tumor type (Q1–3 versus Q4). The proportions of TMB^hi^CD8^low^ and NeoAg^hi^CD8^low^ tumors were compared between expression strata using two-sided Fisher’s exact tests. Effect sizes are reported as odds ratios with corresponding *P* values.

### Robustness across CD8 inference methods

To assess robustness across immune deconvolution methods, CD8⁺ T-cell abundance was estimated using multiple independent approaches, including TIMER^60^, CIBERSORT^55^, and direct RNA-based readouts (*CD8A* and *CD8B* expression). For each method and tumor type, odds ratios for the association between high *NECTIN4* expression (Q4 versus Q1–3) and the TMB^hi^CD8^low^ or NeoAg^hi^CD8^low^ phenotype were calculated. Common odds ratios were estimated using Cochran–Mantel–Haenszel (CMH) models across tumor types. Statistical significance was assessed using two-sided tests.

### Association analyses of *NECTIN4* with immune infiltrates and gene signatures in cancers

Normalized RNA-seq TPM files of TCGA pan-cancer data were obtained from UCSC Xena Browser^61^. For immune cell deconvolution, bulk RNA-sequencing data (TPM) from four TCGA cancer types with high *NECTIN4* expression were analyzed—bladder cancer (BLCA), breast cancer (BRCA), lung adenocarcinoma (LUAD), and lung squamous cell carcinoma (LUSC). Immune cell fractions were inferred using three complementary computational approaches: CIBERSORT^55^, xCell^62^ and TIMER^60^. Similarly, interferon-γ (IFN-γ) response scores were obtained from previously published pan-cancer immune analyses^4^ and integrated with *NECTIN4* expression for downstream correlation analyses. To assess epigenetic regulation of *NECTIN4*, DNA methylation levels across the *NECTIN4* promoter-associated CpG probes were obtained from The NCI Genomic Data Commons. Associations between *NECTIN4* methylation and mRNA expression were evaluated using Spearman correlation and further stratified by *NECTIN4* amplification status to distinguish epigenetic from copy-number–driven effects.

## Single-cell and single-nucleus RNA-seq

### Tumor dissociation and nuclei isolation

Fresh tumor specimens were transferred immediately to ice-cold media following resection and finely minced. Samples were enzymatically and mechanically dissociated using tumor-appropriate digestion cocktails (for example, collagenase and DNase I) to generate single-cell suspensions while minimizing stress-induced transcriptional artifacts. Cell suspensions were filtered sequentially through 70 µm and 40 µm strainers, washed in PBS supplemented with 0.04–1% BSA, and assessed for viability using trypan blue or fluorescence-based methods. Only samples with viability ≥70% were processed further. Cell concentrations were adjusted to ∼700–1,200 cells µl⁻¹ prior to immediate loading onto the 10x Genomics Chromium platform. For frozen tumor specimens, single-nucleus RNA-seq (snRNA-seq) was performed. Frozen tissue blocks were thawed on ice and gently homogenized or triturated in chilled nuclei lysis buffer containing Tris-HCl, NaCl, MgCl₂, and a mild detergent to selectively disrupt cell membranes while preserving nuclear integrity. Nuclei were washed in PBS supplemented with BSA and RNase inhibitor, pelleted at low centrifugal force, filtered through a 40 µm strainer, and resuspended to ∼700–1,200 nuclei µl⁻¹. Optional cleanup steps, including debris or myelin removal, were applied for lipid-rich samples. These approaches follow established 10x Genomics protocols and published tumor sc/snRNA-seq workflows^63,64^.

### 10x Genomics library preparation, sequencing, and preprocessing

Two non–muscle-invasive bladder cancer (NMIBC) samples (BC-FTG-1 and BC-FTG-2) were processed using the 10x Genomics Chromium Single Cell 3′ Gene Expression v3 platform. Cell concentrations and viability were assessed using a LunaFX7 fluorescence cell counter. Libraries were prepared according to the manufacturer’s instructions and sequenced on Illumina platforms. snRNA-seq libraries were generated from three bladder cancer samples (BC-FTG-3, BC-FTG-4, BC-FTG-5) using the 10x Genomics Chromium platform and sequenced on a NovaSeq X Plus system. Base calling was performed using Illumina RTA software. 10x single-cell libraries were processed with Cell Ranger v7.1–v7.2 against GRCh38-2020-A to generate feature-barcode matrices. Read alignment was performed using STAR^65^.

## Single-cell and single-nucleus RNA-seq data analysis

### Data processing and integration

All single-cell and single-nucleus analyses were performed in R (v≥4.2) using Seurat with v4 and v5 compatible workflows^66^. Each dataset was processed as an independent Seurat object. Cells were filtered using established quality filters based on (i) the number of detected genes, (ii) total UMI counts, and (iii) the proportion of mitochondrial transcripts; thresholds were defined per dataset to remove low complexity profiles and outliers. After filtering, datasets were merged and integrated for joint analysis. Batch-associated structure was mitigated using Harmony^67^ on PCA embeddings. We then constructed a shared nearest neighbour graph, performed graph-based clustering, and used UMAP for two-dimensional visualization^68^. Cluster-defining markers were identified using Seurat FindAllMarkers, and cell identities were assigned using canonical lineage markers.

### Definition of epithelial compartments

For bladder cancer datasets, epithelial cells were defined using the lineage annotations of public dataset (GEO: GSE169379) and by canonical epithelial marker expression. Tumor intrinsic analyses were restricted to the epithelial compartment, and epithelial only Seurat objects were reused across downstream analyses to ensure identical cell membership across comparisons. For the breast cancer cohort (GEO: GSE176078)^41^, epithelial cells were selected using curated epithelial barcode annotations supplied with the reference atlas. Epithelial identity was further confirmed by expression of canonical markers including *EPCAM, KRT8, KRT18, KRT19, CDH1, MUC1* and *CLDN3*.

### NECTIN4 quantification and stratification

*NECTIN4* (alias *PVRL4*) was quantified in epithelial cells, which were stratified into *NECTIN4⁺* and *NECTIN4⁻* populations based on log-normalized expression and used for all downstream analyses.

### Differential expression and pathway enrichment

Differential expression between *NECTIN4⁺* and *NECTIN4⁻* epithelial cells was performed using Seurat FindMarkers, and ranked gene lists were then evaluated by Gene Ontology and gene set enrichment analyses (GSEA). Multiple testing was controlled using Benjamini Hochberg false discovery rate correction. Targeted immune-related gene panels were evaluated in parallel, including cytokines and chemokines, immune checkpoint genes, and antigen presentation machinery.

### Immune infiltration and epithelial–immune associations

Immune and other cell populations were quantified at the sample level using marker-based classification. For each sample, lineage-positive fractions were computed and related to epithelial *NECTIN4* metrics (mean expression or fraction of *NECTIN4⁺*epithelial cells). Associations were assessed using Spearman rank correlation for continuous comparisons and non-parametric group comparisons for categorical analyses (including quartile-based stratification with Kruskal Wallis tests, where indicated).

### Copy-number inference from single-cell transcriptomes

Large-scale copy-number alterations were inferred from scRNA-seq data using CopyKAT^40^ applied to raw UMI matrices on a per sample basis. Copy number status at the gene level was summarized from inferred gain and loss burdens and integrated with immune infiltration metrics. Group comparisons were performed using two-sided Wilcoxon rank-sum tests unless otherwise stated.

### Inferring cell-cell communications

Cell–cell communication was inferred from sc and snRNA-seq data using CellChat^69^. CellChat objects were constructed from normalized expression matrices and the curated human ligand-receptor interaction database. We built the intercellular communication networks for *NECTIN4*-positive cells and inferred crosstalk involving other cells. To interrogate candidate NECTIN4 interactors, we augmented the default CellChat ligand-receptor reference with selected biologically supported interactions prior to estimating communication probabilities and network topology.

### Spatial transcriptomics analysis

Public Visium Spatial Gene Expression datasets (fresh-frozen and FFPE, including Visium and Visium HD) were obtained from the 10x Genomics data repository (see reporting summary). For standard Visium datasets, we downloaded the filtered feature barcode matrices together with spatial metadata (tissue images, spot coordinates, and scale factors). For Visium HD datasets, binned expression matrices were analyzed at the selected bin resolution^70^. Data were loaded in Python using Scanpy^71^, and spatial analyses were performed using Squidpy^72^. Tissue inclusion and spatial coordinates were parsed from Space Ranger outputs, and analyses were restricted to spots or bins annotated as within tissue. Image registration and signal distributions were inspected using Loupe Browser (v9). Spatial autocorrelation of *NECTIN4* and immune markers (CD8A, where available) was assessed using Moran’s I computed on spatial neighbor graphs. *NECTIN4* expression was quantified per spot from normalized values; spots with detectable *NECTIN4* expression (>0) were considered *NECTIN4*-positive, and *NECTIN4*-high regions were defined as the top 20% of *NECTIN4*-expressing spots within each sample. Spatial co-localization between *NECTIN4*-enriched and immune-enriched regions was evaluated by neighborhood enrichment with permutation-based Z scores (500 to 1,000 permutations), where positive values indicate spatial attraction and negative values indicate spatial separation. For distance-based analyses, Euclidean distances from each spot or bin to the nearest *NECTIN4* high spot or bin were computed, converting pixel coordinates to microns using Space Ranger scale factors when available. Relationships between immune signature scores and distance to *NECTIN4* high regions were assessed using Spearman rank correlation, and distance stratified summaries were generated for visualization.

### Analysis of RNA-seq and clinical outcomes in immunotherapy cohorts

Clinical and matched transcriptomic data for immunotherapy treated cohorts were obtained from the Cancer Immunology Data Engine^73^. Overall survival data for *NECTIN4* analyses were available for 32 immunotherapy-treated patient cohorts (3,579 patients), and progression-free survival data for 30 cohorts (2,889 patients). Associations between gene expression and survival outcomes were assessed using Cox proportional hazards regression with Wald test p values.

### Somatic variant calling and annotation of whole-genome sequencing data (WGS)

WGS data from TCGA bladder cancer samples were processed using a DRAGEN (v4.2.9) somatic variant-calling workflow. Tumor–normal paired BAM files were obtained from the Genomic Data Commons (GDC) and organized by batch. Somatic variant calling was performed with the Illumina DRAGEN pipeline^74^, producing filtered somatic VCF files for each tumor–normal pair against the hg38 reference genome. In addition to DRAGEN, somatic single nucleotide variants (SNVs) and indels were called using Strelka2 and GATK Mutect2 and separate as well as combined call sets were retained for downstream analyses. Somatic variants were annotated using ANNOVAR^75^ with hg38 gene models. For each sample, VCF files were annotated against multiple reference databases, including RefGene, ClinVar^76^ (2023-04-16 release), dbNSFP v4.2a, COSMIC v92 coding mutations^77^ and dbSNP build 150. Variant-level quality control retained only those variants with FILTER = PASS in the originating VCF. Tumor reference and alternate allele counts were extracted directly from VCF genotype fields (AD, DP, AF), with approximate counts inferred from total depth and allele fraction when explicit allele depth values were unavailable. Variants with total tumor depth ≤10 reads were excluded. To minimize false positives arising from poorly mappable regions, only variants within predefined high-confidence callable regions were retained for analysis.

### Mutation clonality and timing relative to amplification

For NECTIN4-amplified samples, somatic variants were identified using Illumina DRAGEN (v4.2.9). Germline variants were filtered using matched blood normal samples. For each tumor, variant allele depths (AD) and total sequencing depth (DP) were extracted from the VCF fields to compute the variant allele frequency (VAF). For each mutation, the locus-specific total copy number (CN_total) was obtained from segment-level.cns files, and tumor purity (p) was estimated using ABSOLUTE. To determine the clonal status of each mutation, we calculated the Cancer Cell Fraction (CCF), representing the proportion of cancer cells carrying the variant^78^. The CCF was estimated by adjusting the observed VAF for tumor purity and local copy number using the following equation:

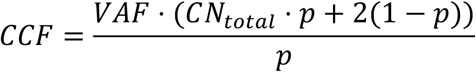

Mutations were classified as clonal-like if CCF ≥ 0.85, subclonal-like if CCF ≤ 0.50 and Intermediate otherwise. For mutations located within NECTIN4 amplicons, multiplicity inference was performed to determine the number of mutated alleles (m). For each possible value of m, the expected VAF was modeled as:

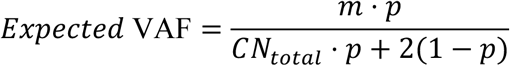

Multiplicity values ranging from m = 1 to m = CN_total were evaluated, and the value minimizing the difference between the observed and expected VAF was selected as the best-fit multiplicity.

Evolutionary timing was inferred based on mutation multiplicity relative to local copy number, following established approaches for timing mutations with respect to copy-number gains^79^. Mutations with m ≥ 4 were classified as pre-amplification, suggesting that the mutation occurred on the ancestral allele prior to the amplification event (or circle formation in ecDNA contexts). Mutations with m = 1 were classified as post-amplification, indicating that the mutation occurred on a single copy after the region had already undergone amplification. Variants with m = 2 or 3 were categorized as uncertain. All computational analyses were performed using Python (v3.12.2) with the Pandas library, and R (v4.2) for statistical visualization.

### Mutational signature deconvolution in BLCA WGS data

Somatic single-nucleotide variants (SNVs) identified from DRAGEN-processed TCGA BLCA whole-genome sequencing (WGS) data were used to construct mutational profiles and perform mutational signature deconvolution. For each sample, somatic mutations were represented as individual Mutation Annotation Format (MAF) files and processed independently using SigProfilerMatrixGenerator (R package)^80,81^. Mutation count matrices were generated in the 96-trinucleotide (SBS96) context using the GRCh38 reference genome, with exome normalization disabled. Each sample was processed separately, without cross-sample aggregation, to ensure that mutational profiles reflected sample-specific processes. Mutational signature attribution was performed using the COSMIC v3.4 SBS reference signature set (https://cancer.sanger.ac.uk/signatures/) and the cosmic fit algorithm implemented in SigProfilerAssignment. Analyses were conducted in matrix input mode with SBS96 context, without collapsing mutation types beyond the standard trinucleotide classification. Given the whole-genome origin of the data, the exome parameter was set to FALSE to apply appropriate normalization and model fitting. All COSMIC SBS signatures were considered, and no prior filtering of signature subsets was applied. Mutation signature profile of bladder cancers from WGS is provided in **Supplementary Data Table 1**.

### Association of recurrent copy-number alterations with mutational signatures

To identify mutational processes associated with recurrent focal copy-number alterations (CNAs), genome-wide CNA profiles were integrated with single-base substitution (SBS) mutational signatures across TCGA bladder cancer samples. Recurrent focal amplifications and deletions were defined using GISTIC calls and restricted to deep deletions or deep amplifications present in ≥10% of tumors. Gene-level focal CNA events were evaluated for association with SBS mutational signatures derived from whole-genome sequencing, focusing on signatures detectable in a sufficient number of samples (present in ≥10% of tumors). Given the large number of candidate SBS mutational signatures, we implemented a two-stage analytical framework combining machine learning–based feature prioritization with statistical hypothesis testing. In the first stage, SBS signature activities were log₂-transformed [log₂(x + 1)] and screened using Random Forest (RF) and XGBoost classifiers trained independently for each focal gene CNA. Feature importance was quantified using Gini impurity (RF) and gain (XGBoost)^82–84^. For each focal CNA, only SBS signatures ranking within the top 20% of importance scores in either model were retained as candidate predictors. This machine-learning step was used exclusively to reduce dimensionality and the multiple-testing burden; importance scores were not used for statistical inference. In the second stage, each retained SBS signature was tested individually for association with CNA status using univariate logistic regression, with CNA presence as the dependent variable and SBS activity as the predictor. Multiple testing was controlled within each focal CNA using Holm’s correction. Using this framework, recurrent focal amplifications on chromosome 1q23—including the *NECTIN4* amplicon and neighboring co-amplified genes—showed strong and consistent associations with APOBEC-associated mutational signatures SBS2 and SBS13.

### JaBbA genome graph reconstruction

Genome-wide junction-balanced genome graphs were reconstructed from TCGA bladder tumor–normal WGS pairs using JaBbA^35^. For each sample, aligned tumor and matched-normal BAM/BAI files were used as input. Candidate rearrangement junctions were obtained with the LOGAN Nextflow pipeline (GRIDSS; hg38), yielding breakpoint-resolved structural-variant calls. JaBbA integrated junction calls with read-depth signals to infer integer copy number on a junction-balanced genome graph. Structural events—including simple classes (deletions, duplications, inversions and translocations) and complex patterns (templated insertion chains, quasi-reciprocal pairs, rigma, pyrgo, tyfonas, breakage–fusion–bridge cycles and double minutes)—were annotated using gGnome, following published workflows^35^.

### DNA isolation, library preparation and Oxford Nanopore sequencing

High–molecular-weight DNA was extracted from tumor tissue and matched peripheral blood using the Nanobind CBB Big DNA Kit (PacBio; 102-301-900). Whole-genome libraries were generated from eight bladder tumors, including five tumor–normal pairs, using the ligation kit SQK-LSK114 and FLO-PRO114M flow cells, and sequenced on a PromethION (run ID: PCA100194). Per-sample yields ranged from 7.23 to 51.41 million pass reads, with median read Q-scores of 16.5–18.5. Basecalling was performed with Dorado v4.3.0 (HAC model) with 5-methylcytosine modification calling enabled.

### Long-read whole-genome sequencing analysis and detection of complex rearrangements

Reads were aligned to the GRCh38 genome using minimap2 (v2.24)^85^ with ‘-k 17-t 28-K 1G - y’ parameters. Clair3 (v1.0.10)^86^, followed by longphase (v1.7.3)^87^, was used to generate phased SNV calls from normal samples for tumor/normal pairs and from tumor samples for the remainder. Whatshap (v2.3) (https://doi.org/10.1101/085050) was used to haplotag both tumor and normal samples. Tumor–normal paired WGS data from five and three tumor only were used for comprehensive characterization of copy-number alterations (CNAs). Somatic structural variants including deletions, duplications, inversions, translocations and complex rearrangements, were detected using Severus^88^. Severus analyzes long-read alignment patterns to generate haplotype-aware base-pair–resolution junction calls. Genome-wide somatic CNAs were inferred using Wakhan^89^ with Severus breakpoints (commit id: 885c7e2), which estimates allele-aware copy-number profiles from sequencing coverage and allelic-imbalance signals. Wakhan produces segmented copy-number profiles across the genome, enabling robust detection of large-scale aneuploidy, focal amplifications and deletions. To resolve focal amplification architecture and distinguish linear amplicons from circular ecDNA-like structures, we applied CoRAL (Complete Reconstruction of Amplifications with Long reads)^90^ to the derived SV and copy-number callsets.

### Breakpoint-based mutational signature and motif analysis

Segment-level copy-number profiles for TCGA bladder tumors were obtained from CNVkit and comprised non-overlapping genomic segments with integer copy-number states. Segments were ordered by genomic coordinate, and breakpoints were defined at boundaries where integer copy number changed between adjacent segments

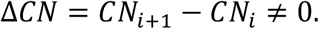

Breakpoints were annotated with chromosome, coordinate, left and right copy-number states, and ΔCN. To limit over-segmentation artefacts, only ≥1 kb segments were used. Local sequence context was extracted from GRCh38 using BSgenome.Hsapiens.UCSC.hg38. For each breakpoint at position *p*, we recorded the trinucleotide context (p−1 to p+1) and enumerated all overlapping trinucleotides within a ±5 bp window (p−5 to p+5). Although COSMIC single-base substitution (SBS) signatures are conventionally defined from SNVs, we used the COSMIC v3 SBS framework^77^ to quantify enrichment of SBS-like trinucleotide contexts at copy-number breakpoints. Two assignment schemes were applied: (i) APOBEC motifs (TCW) were restricted to SBS2/SBS13, with non-APOBEC contexts excluded from SBS2/SBS13; and (ii) remaining contexts were assigned using COSMIC v3 trinucleotide weights. The same procedure was applied to breakpoint sets derived from long-read WGS (Severus)^88^. Signature attribution was performed using both hard and soft assignments. In hard assignment, each breakpoint context was mapped to the single SBS signature with the highest COSMIC weight. In soft assignment, COSMIC weights were normalized to sum to one across compatible signatures for a given context, and breakpoints contributed fractionally to each signature. Per-sample breakpoint-associated signature exposures were obtained by aggregating assignments across all breakpoints.

### Association of APOBEC3 genes and germline variation with *NECTIN4* amplification

To examine relationships between *NECTIN4* amplification and expression of APOBEC3A and APOBEC3B, group-wise comparisons were performed using Wilcoxon rank-sum tests. Associations between germline genetic variation at the *APOBEC3* locus and *NECTIN4* amplification status were evaluated using multivariable regression models incorporating genotypes at rs17000526 (for BLCA) and adjusting for age, sex, and race. Germline genotype data are classified as controlled-access data and were obtained through dbGaP (project 35035).

### Cell culture

Human embryonic kidney 293FT cells (R70007) were purchased from Thermo Fisher Scientific (Waltham, MA, USA). Human bladder cancer cell lines HT-1376 (CRL-1472) and T24 (HTB-4) were obtained from the American Type Culture Collection (ATCC; Manassas, VA, USA). Murine bladder cancer MB49 cells (SCC148) were obtained from Sigma–Aldrich. Commercially sourced cell lines were authenticated by the supplier. All cell lines were routinely tested for mycoplasma contamination. Cells were maintained at 37 °C in a humidified incubator with 5% CO₂. HT-1376 cells were cultured in Eagle’s minimum essential medium (EMEM; ATCC 30-2003) supplemented with 10% fetal bovine serum (FBS). T24 cells were cultured in McCoy’s 5A medium (ATCC 30-2007) supplemented with 10% FBS. 293FT cells were cultured in high-glucose Dulbecco’s modified Eagle medium (DMEM) supplemented with 10% FBS. MB49 cells were cultured in RPMI-1640 medium supplemented with 10% FBS.

### Lentivirus production and infection

Lentiviral particles were produced in 293FT cells by transient co-transfection of a transfer plasmid (7 µg; NECTIN4 human eGFP-tagged Lenti ORF clone, OriGene, RC203431L4) or control vector (7 µg; pLenti-C-mGFP-P2A-Puro, Addgene, 19319), together with the packaging plasmid psPAX2 (6.5 µg; Addgene, 12260) and the envelope plasmid pCMV-VSV-G (1.5 µg; Addgene, 8454). Transfections were performed in 10-cm dishes at approximately 80% confluence using Lipofectamine 3000 (Thermo Fisher Scientific), according to the manufacturer’s instructions. Twelve hours after transfection, the medium was replaced with complete DMEM. Viral supernatants were collected 48 h post-transfection, clarified by centrifugation (3,000 rpm, 10 min, 4 °C), and passed through a 0.45-µm filter before use. For lentiviral transduction, target cells were infected at a multiplicity of infection (MOI) of <1 in the presence of polybrene (10 µg ml⁻¹). Briefly, 4 × 10⁵ cells in suspension (1 ml) were mixed with 1 ml of virus-containing supernatant and subjected to spinoculation (931 g, 45 min, 30 °C), followed by overnight incubation at 37 °C. Puromycin selection (2 µg ml⁻¹) was initiated 48 h after infection for T24 cells and maintained for 4–5 days until all non-transduced control cells were eliminated. Successful NECTIN4–eGFP overexpression was confirmed by immunoblotting and fluorescence microscopy. Vector maps for non-Addgene plasmids are shown in **Supplementary Data Fig. 3**.

### Generation of stable NECTIN4–overexpressing cell lines

Stable overexpression of NECTIN4 was achieved using a commercially available NECTIN4 lentiviral particle (BPS Bioscience, cat. no. 78712). An expression negative control lentivirus (BPS Bioscience, cat. no. 82212-P) was used to generate control cell lines. HT-1376 and T24 cells were seeded in 6-well plates at approximately 40–50% confluence and allowed to adhere overnight. The following day, cells were transduced with lentivirus at a multiplicity of infection (MOI) of 0.5–1 in the presence of 8 µg ml⁻¹ polybrene (MilliporeSigma) to enhance infection efficiency. Cells were incubated with the virus-containing medium for 24 h under standard culture conditions (37 °C, 5% CO₂). After 24 h, the viral supernatant was replaced with fresh growth medium, and cells were cultured for an additional 48–72 h to allow transgene expression.

Stable populations were selected with puromycin (2 µg ml⁻¹) for 7–10 days, with medium replaced every 2–3 days. Stable NECTIN4 overexpression was confirmed by immunoblotting using an anti–NECTIN4 antibody (Abcam, ab192033). Puromycin-resistant NECTIN4-expressing populations were expanded and maintained in puromycin-containing growth medium. Similarly, MB49 monoclonal cell lines were generated using a human NECTIN4 untagged expression plasmid or vector-only control.

### Cell growth assay

T24 cells stably expressing NECTIN4–eGFP were seeded in 96-well tissue culture plates at a density of 4 × 10⁴ cells per well in complete growth medium and allowed to adhere overnight. Cell growth was monitored using an IncuCyte live-cell imaging system (Essen BioScience, Sartorius). Phase-contrast images were acquired every 6 h using a 4× objective. Percent confluence was quantified using IncuCyte analysis software and normalized to the first time point. For longer-term assays, medium was replaced every 2–3 days.

### siRNA-mediated knockdown of select genes

HT-1376 cells were seeded at 3 × 10^5^ cells per well in 6-well plates. At ∼60% confluence the following day, cells were transfected with siRNAs targeting *NECTIN4* (Thermo Fisher Scientific, s37691; antisense 5′-UCAUAUUUCUGGGUCAUCUGC-3′), *DDR1* (Thermo Fisher Scientific, s2298; antisense 5′-AGACUAACCAGAUCUUGAGGG-3′), *PTPN11* (SHP2) (Qiagen, Hs_PTPN11_5; antisense 5′-AGAUGUCCAUGAAACCACCUUdTdT-3′) or a non-targeting control siRNA (Thermo Fisher Scientific, 4390843) at a final concentration of 30 nM. Transfections were performed using Lipofectamine RNAiMAX (Thermo Fisher Scientific) in Opti-MEM Reduced Serum Medium according to the manufacturer’s instructions. Briefly, siRNA and RNAiMAX were diluted separately in Opti-MEM, combined and incubated for 15–20 min at room temperature before addition to cells in antibiotic-free medium. Cells were incubated for 48 h at 37 °C in 5% CO₂ to achieve knockdown. Protein lysates were then prepared for immunoblotting.

### qPCR analysis of cytokine, antigen-presentation and immune-checkpoint gene panel

Total RNA was extracted using the RNeasy Kit (Qiagen) and quantified by spectrophotometry. Complementary DNA (cDNA) was synthesized from 1 µg of total RNA using the RT² First Strand Kit (Qiagen) according to the manufacturer’s instructions, with genomic DNA elimination performed before reverse transcription.

Gene expression profiling was performed using a customized RT² Profiler PCR Array (Qiagen) configured in a 384-well plate format, with each array including HPRT1 for normalization. PCR amplification was performed on a 384-well-compatible real-time PCR system (CFX Opus 384, Bio-Rad) using RT² SYBR Green qPCR Mastermix (Qiagen) under the manufacturer-recommended thermal cycling conditions: 95 °C for 10 min, followed by 40 cycles of 95 °C for 15 s and 60 °C for 1 min. Melt-curve analysis was performed at the end of each run to confirm amplification specificity. Three independent biological replicates were analyzed for each experimental condition.

Raw Ct values were normalized to HPRT1 using the 2^−ΔΔCt^ method, with the control group anchored at a fold change of 1. Differential expression between control and treatment groups was assessed using Welch’s unpaired t-test.

### NECTIN4 protein stimulation

HT-1376 cells were cultured in EMEM supplemented with 10% FBS. Before stimulation, cells were serum-reduced in EMEM containing 1% FBS for 6 h. Recombinant NECTIN4 protein (GenScript, ZZ03823) was reconstituted according to the manufacturer’s instructions and diluted in sterile PBS. Cells were treated with recombinant NECTIN4 at the indicated concentrations for 48 h; vehicle-treated cells served as controls. Cells were washed with ice-cold PBS and lysed in lysis buffer supplemented with protease and phosphatase inhibitors. Lysates were clarified by centrifugation and analyzed by immunoblotting for phosphorylated STAT1. Signals were quantified and normalized to GAPDH.

### Immunoblotting

Cultured cells, either untreated or treated with IFN-γ (Thermo Fisher Scientific, RIFNG100), were lysed in RIPA Lysis and Extraction Buffer (Thermo Scientific, 89901) supplemented with EDTA-free Pierce Protease Inhibitor Tablets (Thermo Scientific, A32959), according to the manufacturer’s instructions. Protein concentrations were determined using the Pierce BCA Protein Assay Kit (Thermo Fisher Scientific). Lysates were mixed with 4× NuPAGE LDS Sample Buffer (Thermo Fisher Scientific) containing 10% β-mercaptoethanol and heated at 95 °C for 10 min. Equal amounts of protein were resolved by SDS–PAGE on NuPAGE Bis-Tris Midi gels (Thermo Fisher Scientific) and transferred to PVDF membranes. Membranes were washed with TBST and blocked for 1 h at room temperature in TBST containing either 5% (w/v) non-fat dry milk or 3% (w/v) BSA. Membranes were incubated overnight at 4 °C with primary antibodies diluted 1:500–1:5,000 in TBST containing 3% BSA, washed three times in TBST (10 min each), and incubated for 1 h at room temperature with HRP-conjugated secondary antibodies (anti-rabbit or anti-mouse IgG; 1:5,000). Bands were visualized using SuperSignal West Pico PLUS (Thermo Scientific, 34580) and quantified by densitometry using ImageJ (NIH). A complete list of antibodies is provided in **Supplementary Data Table 4**.

### Affinity pulldown and mass spectrometry analysis of NECTIN4-interacting proteins

HT-1376 cells were cultured in EMEM supplemented with 10% fetal bovine serum and 1% penicillin–streptomycin under standard conditions (37 °C, 5% CO₂). Cells were seeded into 10-cm dishes in triplicate for each condition at approximately 60% confluence. The following day, cells were transfected with 10 µg NECTIN4–Myc–DDK–tagged expression plasmid (OriGene, RC203431) using Lipofectamine 3000 (Thermo Fisher Scientific), according to the manufacturer’s instructions. As a negative control, cells were transfected in parallel with an equal amount of the pCMV6-Entry mammalian expression vector (OriGene, PS100001).

After 48 h, cells were harvested and lysed for immunoprecipitation using the c-Myc–tagged Protein Mild Purification Kit ver. 2 (MBL International, 3305). Cell pellets were resuspended in lysis buffer (50 mM Tris-HCl pH 7.5, 150 mM NaCl, 5 mM EDTA, 1% Triton X-100) and gently rotated at 4 °C for 30 min. Lysates were clarified by centrifugation at 15,000 × *g* for 10 min at 4 °C. The resulting supernatants were transferred to the kit-supplied spin columns and incubated with anti–c-Myc affinity beads for 1 h at 4 °C with gentle end-over-end rotation. Beads were then washed five times with the supplied wash buffer and finally resuspended in 10 µl 50 mM Tris-HCl to prevent drying prior to mass spectrometry.

Immunoprecipitated proteins were eluted in 5% SDS and processed for on-column tryptic digestion using mini spin columns (Epoch Life Science), as previously described^91^. Briefly, samples were precipitated with methanol/ammonium bicarbonate, loaded onto columns, washed, and digested overnight with trypsin (37 °C). Peptides were eluted with 1% acetic acid, dried, resuspended in 3% acetonitrile/0.1% trifluoroacetic acid, quantified by A205, and equal amounts were injected. Peptides were analyzed by nanoLC–MS/MS on an UltiMate 3000 RSLCnano coupled to an Orbitrap Fusion Lumos (Thermo Scientific) operated in data-independent acquisition (DIA) mode. Peptides were separated on a 75 µm × 15 cm EASY-Spray column using a 0.1% formic acid/water (A) and 0.1% formic acid/acetonitrile (B) gradient (107 min; 300 nL min⁻¹). MS1 spectra were acquired at 120,000 resolution (m/z 420–680), and DIA MS2 spectra were acquired at 60,000 resolution using 60 variable windows (4 m/z) across m/z 430–670 with HCD (27% NCE). Each sample was analyzed in technical duplicate. DIA data were processed in DIA-NN (v2.2.0) using an in silico–predicted spectral library derived from the UniProt human database, with carbamidomethylation (C) as a fixed modification and one missed cleavage allowed. Protein inference was performed at the gene level with RT-dependent cross-run normalization and robust LC quantification; other parameters were left at recommended/default settings. Proteins enriched in NECTIN4 pulldowns relative to empty-vector controls across both pulldown approaches were considered candidate interactors, and DDR1 was prioritized for further investigation.

### Immunoprecipitation

HT-1376 bladder cancer cells were cultured in EMEM supplemented with 10% fetal bovine serum and 1% penicillin–streptomycin at 37 °C in a humidified incubator with 5% CO₂. Cells were seeded in 10-cm dishes at approximately 60% confluence and transfected the following day with 10 µg NECTIN4–Myc–DDK–tagged expression plasmid (OriGene, RC203431) using Lipofectamine 3000 (Thermo Fisher Scientific), according to the manufacturer’s instructions. Cells transfected with empty vector (pCMV6-Entry; OriGene, PS100001) served as negative controls. Cells were harvested 48 h post-transfection and lysed in buffer containing 50 mM Tris-HCl (pH 7.5), 150 mM NaCl, 5 mM EDTA and 1% Triton X-100 for 30 min at 4 °C with gentle agitation. Lysates were clarified by centrifugation at 15,000 × *g* for 10 min at 4 °C. For immunoprecipitation, clarified lysates were incubated with Pierce Anti-DYKDDDDK (FLAG) affinity resin (Thermo Fisher Scientific) for 4 h at 4 °C with end-over-end rotation. The resin was washed five times with lysis buffer, and bound proteins were eluted by boiling the beads in SDS sample buffer. Eluted proteins were resolved by SDS–PAGE and analyzed by immunoblotting using anti–Myc-tag and anti-DDR1 antibodies (see reporting summary).

### Syngeneic murine tumor implantation, antibody treatment and tumor-growth analysis

MB49 bladder cancer cells stably transduced with either a control vector or a human NECTIN4 expression construct were used for syngeneic tumor implantation studies. Female C57BL/6 mice were acclimatized for 1–2 weeks and maintained under specific pathogen-free conditions at the National Cancer Institute animal facility. All procedures were conducted in accordance with approved institutional animal-care guidelines. Cells were cultured under standard conditions to 70–80% confluence, detached, washed twice with sterile PBS and resuspended in serum-free RPMI at 5 × 10⁶ cells ml⁻¹. A total of 5 × 10⁵ cells in 50 µl RPMI was mixed 1:1 with Matrigel (Corning) and injected subcutaneously into the flank of immunocompetent C57BL/6 mice in a final volume of 100 µl. Mice were randomly assigned to experimental groups. For treatment studies, mice bearing parental MB49 or MB49-NECTIN4 tumors were treated with control IgG, anti–PD-1 antibody or a NECTIN4-targeting antibody at 10 mg kg⁻¹ according to the dosing schedule shown in the corresponding figure legend. The antibodies used for in vivo treatment were anti-mouse PD-1 antibody (Bio X Cell, BE0033-2), control IgG (Bio X Cell, BE0091) and anti-human NECTIN4 antibody/enfortumab biosimilar (Bio X Cell, SIM0031). Tumor dimensions were measured longitudinally using calipers, and tumor volume was calculated as:

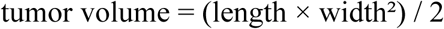

where length represents the longest tumor dimension and width represents the perpendicular dimension. Tumor growth was monitored until the experimental endpoint or until tumors approached institutional humane endpoints.

Tumor-growth analyses were performed using absolute tumor volumes. Because dropout related to tumor burden occurred after day 15 in control groups, the primary longitudinal analysis was restricted to measurements collected through day 15. Tumor-growth kinetics were analyzed using linear mixed-effects models with log₂-transformed tumor volume as the outcome:

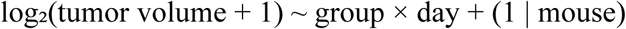

where group, day and their interaction were modeled as fixed effects, and mouse was included as a random intercept to account for repeated measurements within individual animals. Treatment effects on tumor-growth rate were assessed using planned contrasts of group-specific slopes estimated from the mixed-effects model. Specifically, slope contrasts were used to compare MB49-NECTIN4 versus parental MB49 tumors under IgG treatment, anti–PD-1 versus IgG treatment within parental MB49 tumors, anti–PD-1 versus IgG treatment within MB49-NECTIN4 tumors, and NECTIN4-targeting antibody versus IgG treatment within MB49-NECTIN4 tumors. *P* values were derived from mixed-effects model slope contrasts. Endpoint tumor volumes and area under the tumor-growth curve were evaluated as supportive analyses where indicated. Data are shown as mean ± s.e.m. unless otherwise stated. Statistical analyses were performed in R using the lme4, lmerTest and emmeans packages. Investigators were not blinded to group allocation during treatment, but tumor measurements and downstream analyses were performed using consistent criteria across groups.

### Flow-cytometric analysis

For immune-profiling studies, mice were euthanized three weeks after implantation, and tumors were excised for downstream flow-cytometric analysis of immune-cell populations in the tumor microenvironment. Mice that failed to develop tumors or exhibited extensive tumor necrosis before the three-week harvest were excluded from flow-cytometric analyses. Independent experiments were performed on separate dates using independently prepared cells, and gating was performed using an identical strategy across experiments.

Tumor tissues were finely minced into 0.5–1.0-mm fragments and processed into single-cell suspensions using the Tumor Dissociation Kit (130-096-730, Miltenyi Biotec) according to the manufacturer’s instructions. Following mechanical and enzymatic dissociation, cell suspensions were passed through a MACS SmartStrainer (70 µm), and erythrocytes were lysed using ACK lysing buffer. Cells were washed, resuspended in FACS buffer consisting of PBS with 2% FBS and kept on ice before counting and antibody staining.

The antibody panel included CD45–Alexa Fluor 700 (147715, BioLegend) to distinguish hematopoietic cells from stromal and epithelial compartments, and T-cell markers CD3ε–PE (12-0031-82, eBioscience), CD4–Brilliant Violet 605 (100547, BioLegend) and CD8–FITC (100706, BioLegend). Viable cells were identified using Zombie UV viability dye (BioLegend). Compensation controls were prepared for each fluorophore using UltraComp eBeads Compensation Beads (Invitrogen). Multiparameter flow-cytometry data were acquired on a BD FACSymphony A5 or BD LSRFortessa SORP cytometer and analyzed using FlowJo v10 software (BD Biosciences).

### Super-resolution immunofluorescence microscopy

A total of 35,000 cells were seeded onto 12 mm poly-D-lysine–coated glass coverslips 24 h before fixation. Cells were washed with ice-cold PBS and fixed in 4% paraformaldehyde in PBS, followed by three PBS washes. Wheat germ agglutinin was applied at 2 µg ml⁻¹ for 20 min to label cell boundaries, then removed by three PBS washes. Coverslips were blocked for 30 min in PBS containing 1% BSA and glycine (22.52 mg ml⁻¹). Primary antibodies were diluted in 1% BSA in PBS and incubated overnight at 4 °C with gentle agitation. After three PBS washes, coverslips were incubated with secondary antibodies diluted in 1% BSA in PBS for 1 h with agitation. Nuclei were counterstained with DAPI for 5 min. Coverslips were washed four times in PBS and mounted using ProLong Glass (Invitrogen). Z-stacks were acquired on a Nikon SoRa spinning-disk microscope (CCR Microscopy Core) using a 60× Apo TIRF oil objective (NA 1.49) and a Photometrics BSI sCMOS camera. Images were processed in NIS-Elements AR (v6.10.02) using the denoise.ai module.

## Statistics, and reproducibility

All statistical analyses were performed using R (v4.x) with ggplot2 and related packages. Computational analyses were performed on the NIH Biowulf high-performance computing cluster. Specific statistical tests, sample sizes (*n*), and multiple-testing correction procedures are described in the corresponding Methods sections and Fig. legends. All statistical tests were two-sided unless otherwise specified. No statistical methods were used to predetermine sample size.

Five normal urothelium samples were excluded from downstream analyses and were used only for copy-number profiling and structural variant detection in cases with reliable ploidy and purity estimates. Nine TCGA tumor samples were excluded from DRAGEN somatic mutation calling and SigProfilerAssignment analyses because of data-quality considerations. TCGA matched BLCA normal samples were excluded from all analyses except somatic mutation calling and copy-number modeling using JaBbA.

Mouse tumor immune infiltration assays assessing the effects of NECTIN4 overexpression were performed independently two times. RNA-sequencing experiments of enfortumab vedotin–treated cells were conducted with three independent biological replicates per condition. Mass spectrometry analyses of pulldown samples were performed in triplicates. siRNA-mediated *NECTIN4* knockdown and overexpression experiments were conducted with a minimum of three independent biological replicates and typically exceeded three replicates during repeated validation and antibody screening for western blot analyses. Independent biological replicates are indicated in the figures and their legends.

## Data records and availability

DNA and RNA sequencing data from patient samples have been deposited in the Database of Genotypes and Phenotypes (dbGaP) under accession number phs004506.v1. Processed single-cell/nucleus RNA-seq data with gene–cell count matrices have been deposited in the Gene Expression Omnibus (GEO) under accession number GSE319003. Proteomics mass spectrometry data have been deposited in the ProteomeXchange Consortium via the PRIDE repository under accession number PXD073805. Plasmid information is provided in **Supplementary Data Fig. 3**. Figure source data will be provided with the journal submission and are available from the corresponding author upon reasonable request.

## Code availability

The following custom, open-source code and databases/tools were used in this article. Genomic and transcriptomic analyses used JaBbA (v1.1) (https://github.com/mskilab/JaBbA), gGnome (commit c390d80) (https://github.com/mskilab/gGnome), AmpliconArchitect (https://github.com/virajbdeshpande/AmpliconArchitect), FishHook (commit 06e3927) (https://github.com/mskilab/fishHook), SigProfilerAssignment (https://github.com/SigProfilerSuite/SigProfilerAssignment), GATK (v4.1.0) (https://github.com/broadinstitute/gatk), Ensembl (v93) (https://www.ensembl.org), COSMIC (v3.4) (https://cancer.sanger.ac.uk), ClinVar (2023-04-16 release) (https://www.ncbi.nlm.nih.gov/clinvar/), dbSNP (v150) (https://www.ncbi.nlm.nih.gov/snp/), Scanpy (v1.9.6) (https://github.com/scverse/scanpy), CycleViz (v0.1.5) (https://github.com/AmpliconSuite/CycleViz), CellRanger (v7.1.0) (https://github.com/10XGenomics/cellranger), ANNOVAR (https://github.com/WGLab/doc-ANNOVAR), CIDE (https://cide.ccr.cancer.gov), Severus (v1.2) (https://github.com/KolmogorovLab/Severus). Additional tools included CopyKAT (https://github.com/navinlabcode/copykat), MsigDB (https://www.gsea-msigdb.org/gsea/msigdb/collections.jsp), Mutect2 (https://doi.org/10.1101/861054), DRAGEN (https://doi.org/10.1038/s41587-024-02382-1), CellChat (https://github.com/sqjin/CellChat), CNVkit (https://github.com/etal/cnvkit), CoRAL (DOI: 10.1101/gr.279131.124), Wakhan (https://github.com/KolmogorovLab/Wakhan), Squidpy (https://github.com/scverse/squidpy), CIBERSORTX (https://cibersortx.stanford.edu), and TIMER3 (https://compbio.cn/timer3/). Figures and image processing used BioRender.com, Fiji/ImageJ (v154f), GraphPad Prism (v10.2.0), SnapGene, FlowJo (v10), IncuCyte (Essen BioScience, Sartorius), and NIS-Elements AR (v6.10.02; denoise.ai). CIDE data were downloaded before October 30, 2025. Custom code for the residual-based regression framework and breakpoint/mutational-signature analyses is available on GitHub (https://github.com/roufbanday/WGS_Bladder_cancer_Manuscript_Figures) and archived on Zenodo.

## Acknowledgements

This work was supported in part by the Intramural Research Program of the National Institutes of Health (NIH). Contributions by NIH authors were made as part of their official duties as federal employees and are considered works of the United States Government. The findings and conclusions are those of the authors and do not necessarily reflect the views of the NIH or the U.S. Department of Health and Human Services. A.R. Banday, A.B. Apolo, and M. Kolmogorov were supported by the Intramural Research Program of the Center for Cancer Research, National Cancer Institute, National Institutes of Health, through grants 1ZIABC012091, 1ZIABC011351, and 1ZIABC012104, respectively. We thank Dr. Michael Kruhlak, Mr. Langston Lim and Dr. Andy Tran (CCR Microscopy Core) for assistance with immunofluorescence imaging; Dr. Bao Tran, Dr. Yongmei Zhao and Mr. Biraj Shrestha (CCR Frederick Sequencing and Genomics Core) for RNA and DNA sequencing and initial data processing; and Dr. Shafiuddin Siddiqui (CCR/LGI Flow Cytometry Core) for guidance on flow cytometry analytical and sorting experiments. The authors used NIH CHIRP and ChatGPT for proofreading and grammatical refinement of the manuscript text.

## Author contributions

A.R.B., B.L. and D.Y. conceived and designed the study. B.L., A.C., J.R., T.C.A., A.Y.R., W.Y., L.M.J. and S.B. performed experiments. D.Y., E.U., A.G.K., K.B., M.K. and A.R.B. conducted statistical and bioinformatics analyses. R.C., E.B.A.C., V.A.V.R., S.G. and A.B.A. contributed patient care, clinical work and surgical procedures. A.R.B. supervised the research. A.R.B, B.L., and D.Y. wrote the first draft. All authors contributed to revision and editing and approved the final manuscript.

