## Supplementary Figures 1-10 for "APOBEC3-driven neoantigen-rich cancers co-opt 1q23.3 amplification for tumor-intrinsic immune cloaking"

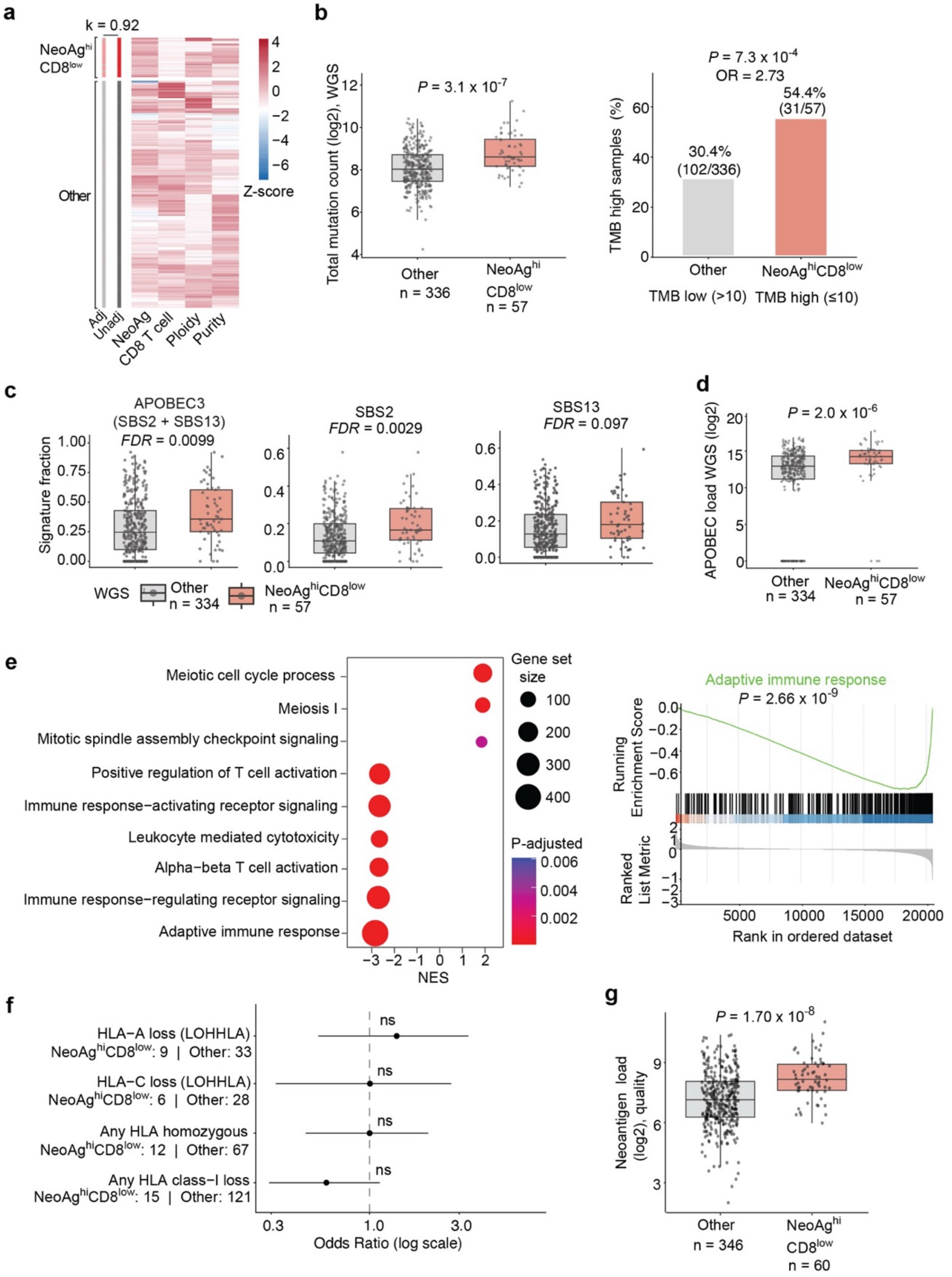

**Supplementary Fig. 1 | Characterization of the NeoAg$^{hi}$CD8$^{low}$ immune–neoantigen-discordant phenotype.**
**a,** Heatmap showing concordance of NeoAg$^{hi}$CD8$^{low}$ tumor classification before and after adjustment for tumor purity and ploidy. Rows represent tumors, and columns show neoantigen load, CD8$^{+}$ T-cell infiltration, ploidy and purity. Cohen's $\kappa = 0.92$ indicates high concordance between unadjusted and adjusted classifications.
**b,** Left, total somatic mutation count from whole-genome sequencing in NeoAg$^{hi}$CD8$^{low}$ tumors and other tumors. *P* value was calculated using a two-sided Wilcoxon rank-sum test. Right, proportion of TMB-high tumors in each group. Numbers indicate sample counts. P value and odds ratio were calculated using Fisher's exact test.
**c,** APOBEC3-associated mutational signature fractions in NeoAg$^{hi}$CD8$^{low}$ tumors and other tumors. The composite APOBEC3 fraction represents the sum of SBS2 and SBS13. Signature fractions were derived from whole-genome sequencing using SigProfiler. *P* values were calculated using two-sided Wilcoxon rank-sum tests with Benjamini–Hochberg correction.
**d,** APOBEC3-associated mutation load from whole-genome sequencing in NeoAg$^{hi}$CD8$^{low}$ tumors and other tumors. P value was calculated using a two-sided Wilcoxon rank-sum test.
**e,** Left, gene set enrichment analysis of differentially expressed genes in NeoAg^hiCD8^low tumors compared with other tumors using GO gene sets. Dot size represents gene set size, and color indicates adjusted P value. Right, GSEA enrichment plot for the Adaptive Immune Response gene set. NES, normalized enrichment score.
**f,** Forest plot showing associations between HLA class I loss features and the NeoAg$^{hi}$CD8$^{low}$ phenotype. HLA loss features included LOHHLA-derived allele-specific loss calls, HLA homozygosity and nonsense mutations affecting *HLA-A*, *HLA-B*, *HLA-C* or *B2M*. Odds ratios are shown on a log scale with 95% confidence intervals. P values were calculated using Fisher's exact tests with Benjamini–Hochberg correction; ns, not significant.
**g,** Predicted neoantigen quality in NeoAg$^{hi}$CD8$^{low}$ tumors versus other tumors. *P* value was calculated using a two-sided Wilcoxon rank-sum test.

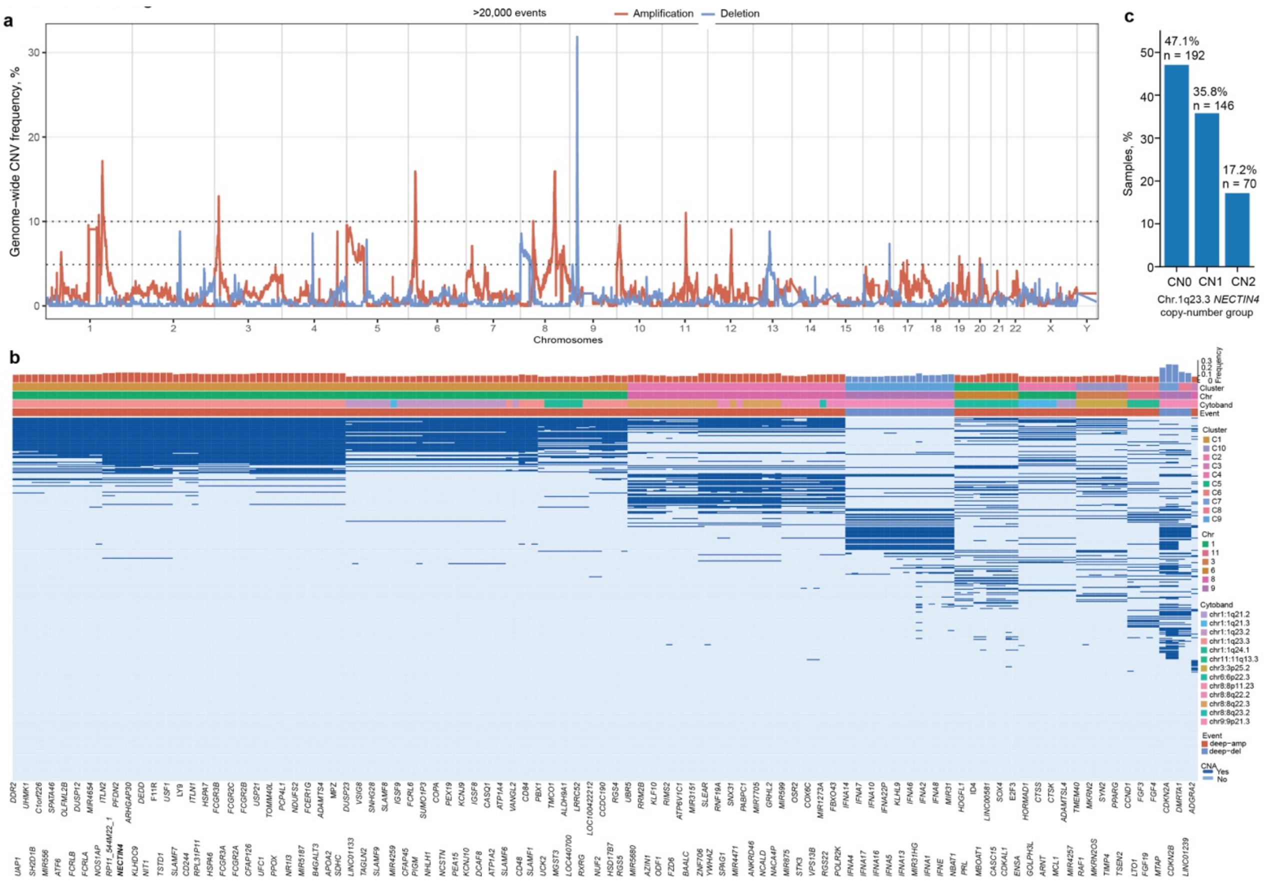


**Supplementary Fig. 2 | Recurrent copy-number alterations in bladder cancer and their co-occurrence as genomic clusters.**

**a,** Manhattan plot of recurrent copy-number alterations in TCGA bladder cancer (BLCA) based on GISTIC calls obtained from cBioPortal. The y axis shows alteration frequency across samples, and the x axis shows genomic position ordered by chromosome and genomic coordinate (GRCh38). Blue peaks denote focal deletions, and red peaks denote focal high-level amplifications. The dashed horizontal line marks the recurrence threshold of alterations present in ≥10% of samples.

**b,** Heatmap depicting clusters of recurrent deep deletions and amplifications present in ≥10% of samples. Hierarchical clustering was performed using CNA co-occurrence rates, with clusters defined at a co-occurrence threshold of ≥75%. Clusters are annotated by chromosome and cytoband, and genes are assigned to their corresponding clusters. Gene names are shown along the x axis, with bars above the heatmap indicating the alteration frequency for each gene. Ten distinct CNA clusters were identified; *NECTIN4* resides within Cluster 1. Gene labels are shown on two staggered lines to improve readability.

**c,** Fraction of TCGA BLCA tumors stratified by *NECTIN4* copy-number status at chromosome 1q23.3. CN0 denotes no copy-number gain, CN1 denotes single-copy gain and CN2 denotes high-level amplification. *NECTIN4* was among the most frequently copy-number-altered genes within the chromosome 1q23.3 cluster.

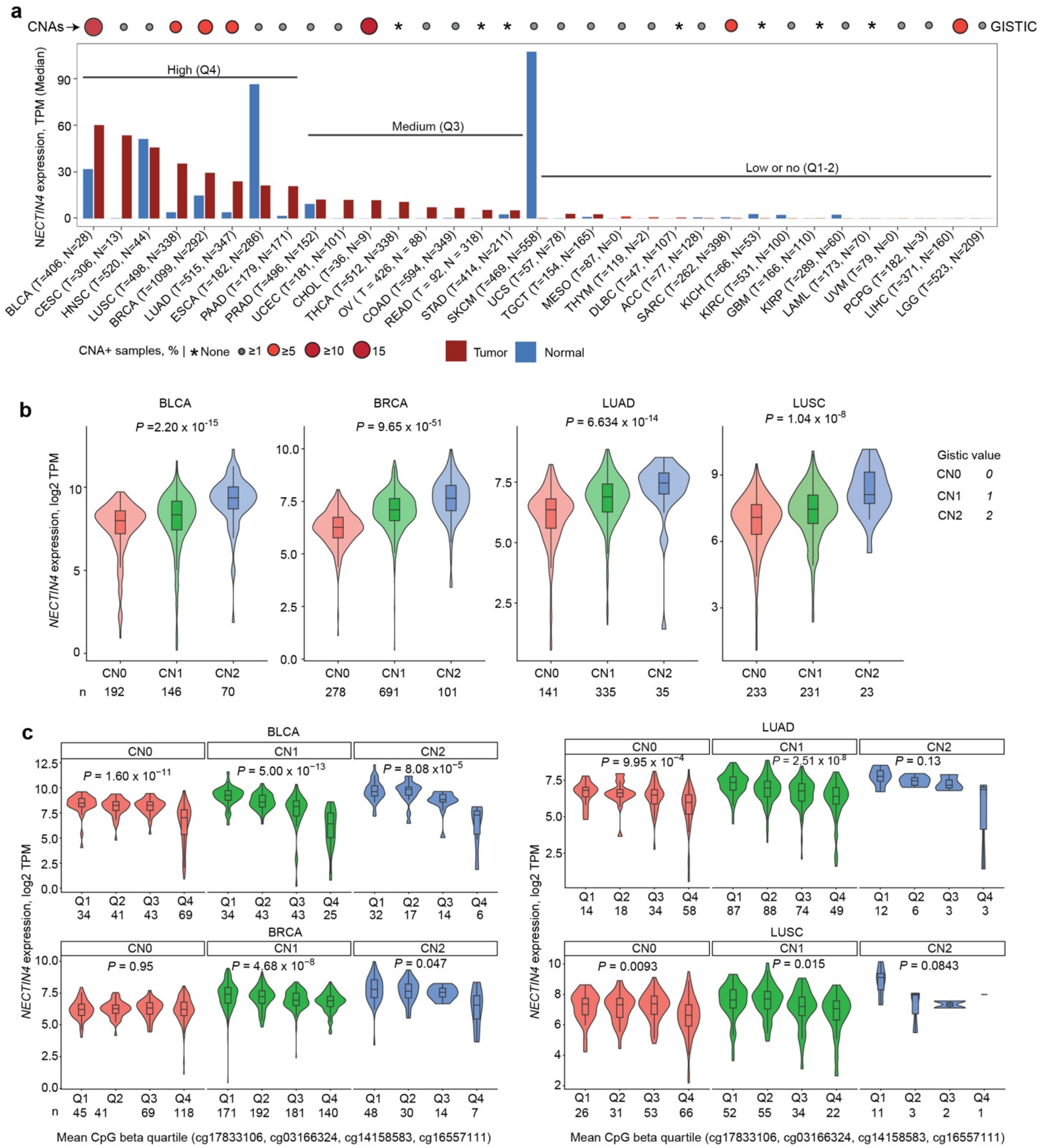


**Supplementary Fig. 3 | Copy-number and epigenetic regulation of *NECTIN4* expression across cancers.**

**a,** Bar plot showing median *NECTIN4* mRNA expression (TPM) across TCGA pan-cancer cohorts in tumor (red) and matched normal tissues (blue). Red circles above each cancer type indicate the frequency of GISTIC2-defined *NECTIN4* amplification events. Tumors are grouped into high (Q4), intermediate (Q2), and low or absent (Q1–Q3) *NECTIN4* expression categories based on cohort-specific quartiles. Sample sizes for each cohort are indicated.

**b,** Violin plots showing *NECTIN4* expression stratified by GISTIC2 copy-number state (CN0: diploid; CN1: low-level gain; CN2: high-level amplification) in BLCA, BRCA, LUAD, and LUSC. Center lines denote medians and interquartile ranges. *P* values were calculated using two-sided Kruskal–Wallis tests.
**c,** *NECTIN4* expression stratified jointly by copy-number state and promoter CpG methylation quartiles (cg17833106, cg03166324, cg14158583, cg16557111) in BLCA, BRCA, LUAD, and LUSC. CpG quartiles were defined within each cohort. *P* values denote associations between methylation quartile and expression within each copy-number category (two-sided Kruskal–Wallis tests). Sample sizes per group are indicated.

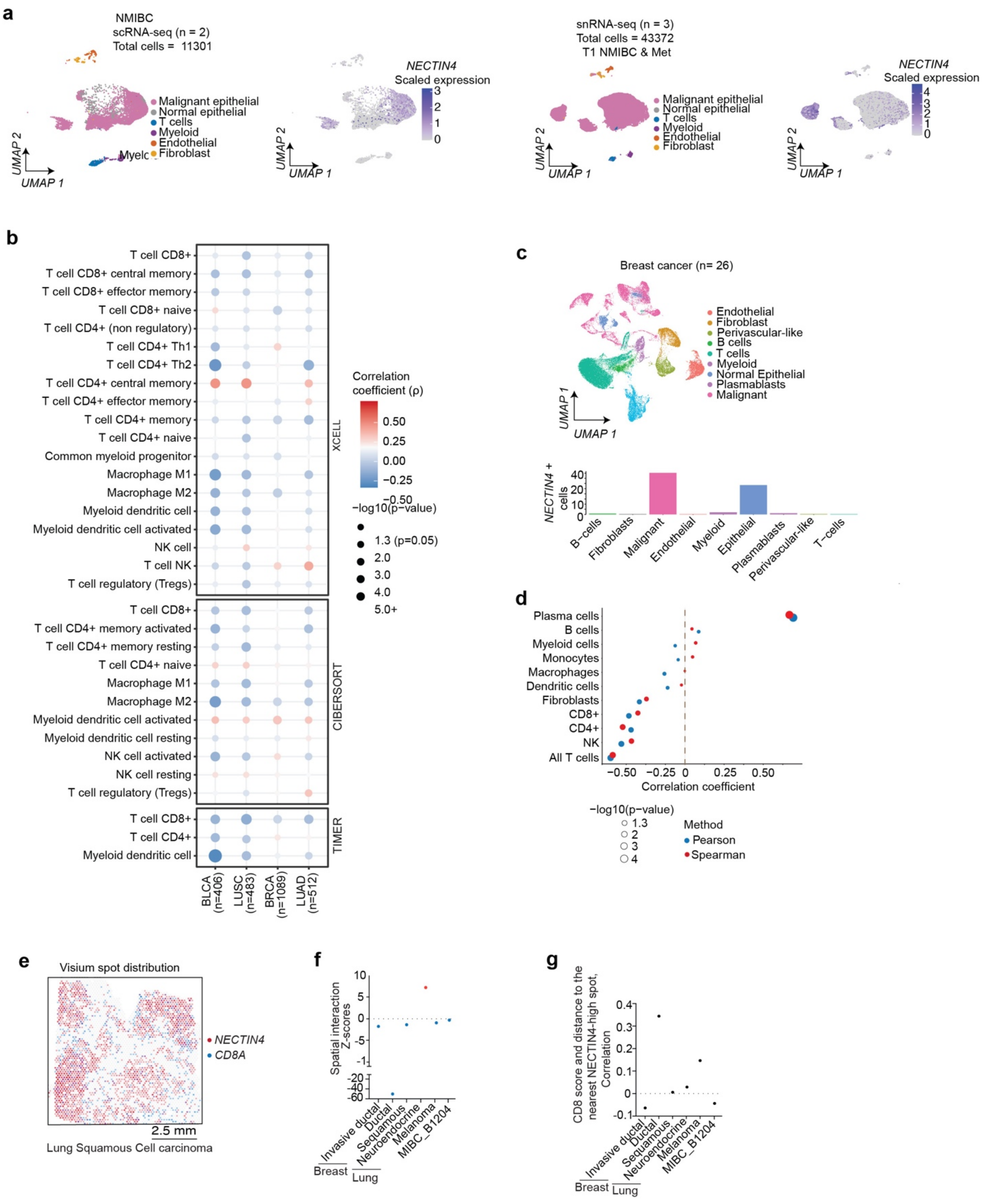


**Supplementary Fig. 4 | *NECTIN4* expression associates with an immune-cold tumor microenvironment across cancers.**

**a,** UMAP visualization of single-cell (scRNA-seq; NMIBC, n = 2) and single-nucleus (snRNA-seq; NMIBC/Met, n = 3) bladder cancer datasets. Left panels are annotated by major cell types. Right panels show *NECTIN4* expression, with colour intensity indicating scaled expression levels.
**b,** Dot plots showing Spearman correlation analyses between *NECTIN4* expression and immune cell populations inferred from TCGA bulk RNA-sequencing data across multiple tumor types using three independent deconvolution methods: xCell, CIBERSORT, and TIMER. Dot colour represents the Spearman correlation coefficient (ρ), and dot size indicates $-\log_{10}(P \text{ value})$.
**c,** Upper panel: UMAP visualization of major cell types from single-cell RNA-sequencing of breast tumors (n = 26). Lower panel: Bar plot showing the fraction of *NECTIN4*$^{+}$ cells across annotated cell populations.
**d,** Associations between *NECTIN4* expression in epithelial cells and immune cell populations in breast cancer, assessed using Pearson (blue) and Spearman (red) correlation analyses. The x axis shows correlation coefficients, and dot size represents $-\log_{10}(\text{P value})$.
**e,** Representative Visium spatial transcriptomics spot map from a lung squamous cell carcinoma sample, with *NECTIN4*-high spots shown in red and *CD8A*-positive spots shown in blue.
**f, g,** Scatter plots show the relationship between *CD8A* expression and the distance to the nearest *NECTIN4*-high spot (f), and corresponding spatial interaction Z scores (g). Across tumor types, in samples with positive Moran's I for a given gene, both analyses indicate spatial exclusion of *CD8A*-positive cells from *NECTIN4*-high tumor regions. Samples with non-positive Moran's I—owing to absent expression, expression in <1% of spots, or non-computable statistics—were excluded.

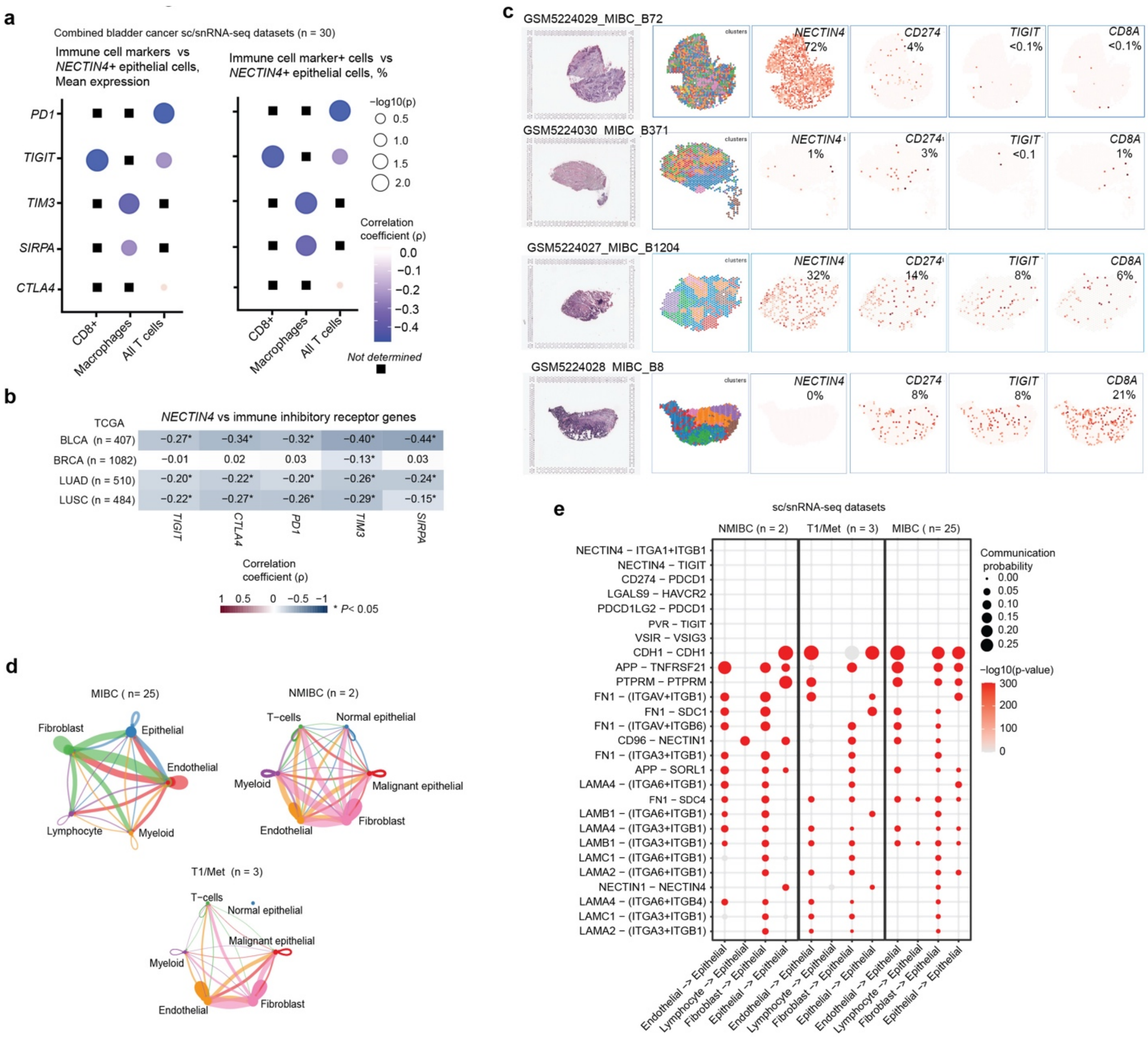


**Supplementary Fig. 5 | *NECTIN4*-expressing malignant cells exhibit limited direct lymphocyte interactions based on single-cell RNA sequencing data.**

**a,** Dot plots showing Spearman's rank correlation analyses between average *NECTIN4* expression in malignant epithelial cells and immune checkpoint receptors expressed on specified immune cell populations in combined single-cell and single-nucleus RNA-sequencing (sc/snRNA-seq) datasets (left). Right, associations between the fraction of *NECTIN4*$^+$ epithelial cells and the fraction of immune cell marker–positive cells across tumors. Dot size represents $-\log_{10}(P$ value), and dot color indicates the Spearman correlation coefficient (ρ).

**b,** Heatmap showing Spearman rank correlations between selected immune checkpoint genes and *NECTIN4* mRNA expression in BLCA, BRCA, LUAD and LUSC. Colors indicate the direction and magnitude of correlation (positive or negative), and correlation coefficients are annotated. Asterisks denote statistically significant associations ($P < 0.05$).

**c,** Spatial layout of tumor samples from muscle-invasive bladder cancer (MIBC), including corresponding H&E-stained sections and spatial expression patterns of *NECTIN4*, PD-L1 (*CD274*), *TIGIT*, and *CD8A*.
**d,** Cell–cell interaction network inferred using the CellChat framework, highlighting interactions between *NECTIN4*-expressing epithelial and endothelial cells and distinct populations of *NECTIN4*-negative cells. Edge thickness indicates the relative strength of inferred interactions.
**e,** Dot plot showing inferred ligand–receptor interactions (y axis) across interacting cell types (x axis), derived from single-cell and single-nucleus RNA sequencing data. The top 20 signals and curated ligand–receptor pairs are shown. Dot size reflects interaction strength, and color denotes relative signaling probability.

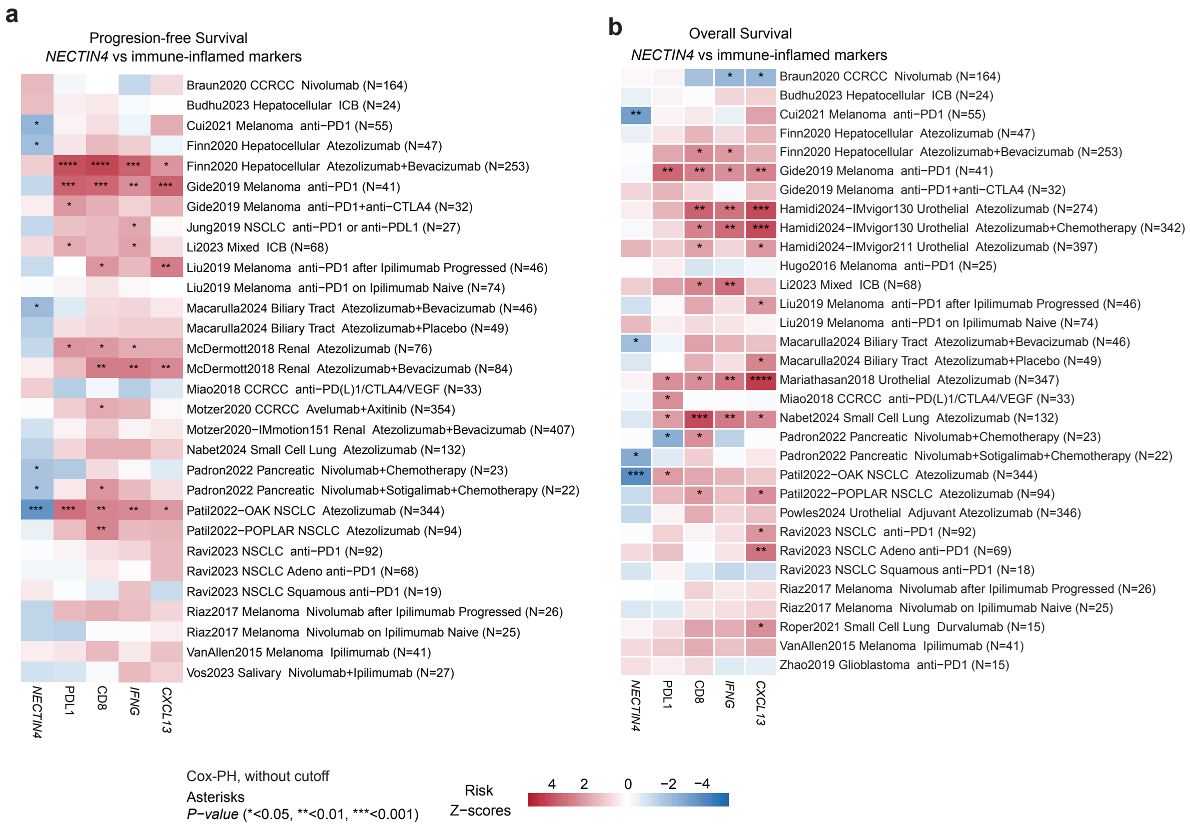


**Supplementary Fig. 6 | Association of NECTIN4 expression with clinical outcomes in immune checkpoint blockade–treated cohorts.**

**a,b,** Heatmaps showing associations between *NECTIN4* mRNA expression and progression-free survival (**a**) or overall survival (**b**) across immune checkpoint blockade–treated cohorts curated from the Cancer Immunology Data Engine (CIDE). Associations for representative immune-inflamed markers or signatures, including PD-L1, CD8, *IFNG* and *CXCL13*, are shown for comparison. Risk Z-scores and P values were derived from Cox proportional hazards regression models. Colors indicate the direction and magnitude of association for higher expression of the indicated marker or signature, with red indicating association with improved outcome and blue indicating association with poorer outcome. Asterisks denote statistical significance ($P < 0.05$, $P < 0.01$ and $P < 0.001$). Higher *NECTIN4* expression was recurrently associated with poorer progression-free and overall survival across multiple cohorts.

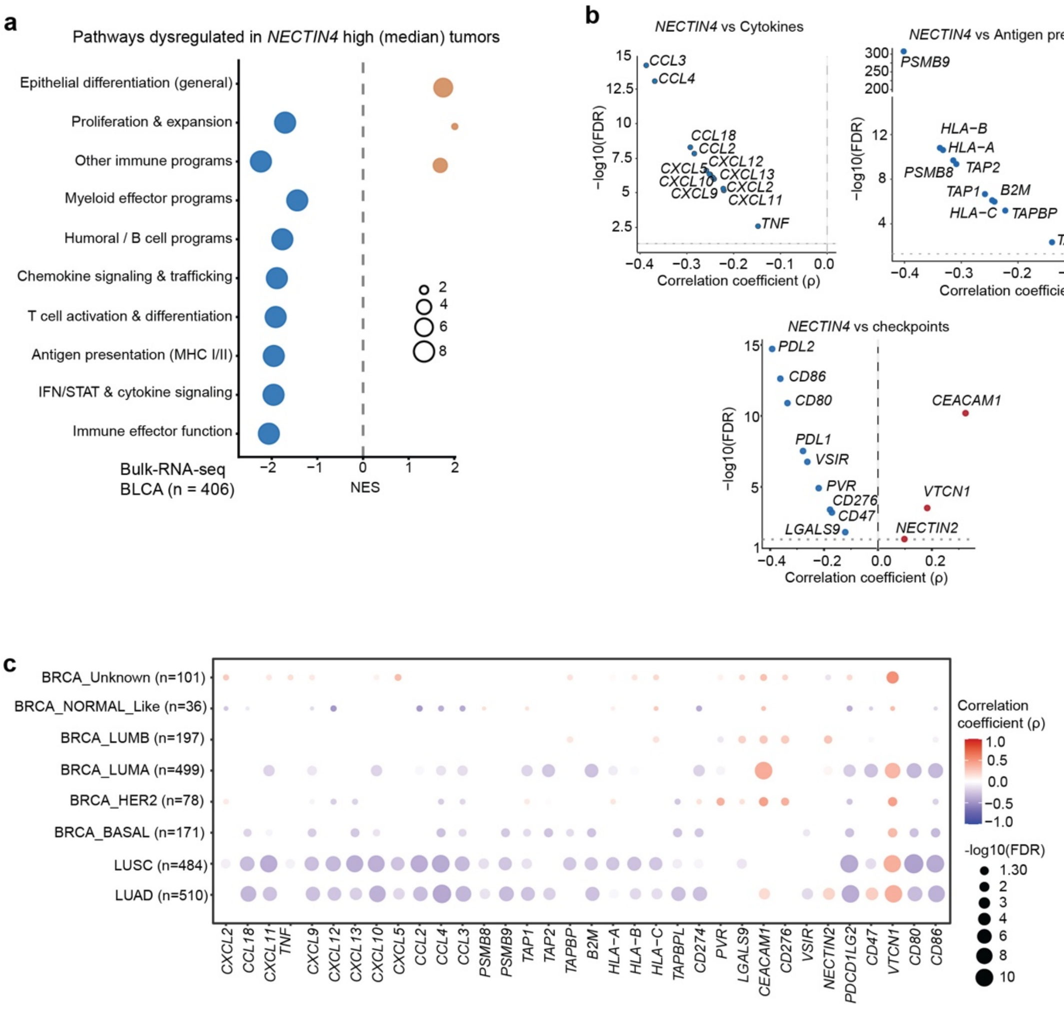


**Supplementary Fig. 7 | NECTIN4 expression is associated with suppression of antigen presentation and immune checkpoint programs in malignant cells.**

**a,** Pathway enrichment analysis in TCGA BLCA (n = 406) comparing *NECTIN4*-high versus *NECTIN4*-low tumors (median split). Dot color represents normalized enrichment score (NES), and dot size indicates $-\log_{10}$(FDR).

**b,** Spearman correlation analyses between *NECTIN4* mRNA expression and immune-related gene modules in TCGA BLCA (n = 406), including cytokines/chemokines, antigen presentation machinery, and immune checkpoint genes. The x axis shows Spearman's rank correlation coefficient (ρ), and the y axis shows $-\log_{10}$(FDR).

**c,** Spearman correlation analyses between *NECTIN4* expression and selected cytokine, antigen presentation, and immune checkpoint genes across tumor types. For breast cancer, results are shown stratified by molecular subtype to illustrate subtype-specific patterns. Dot size represents $-\log_{10}$(FDR), and color indicates Spearman's rank correlation coefficient (ρ).

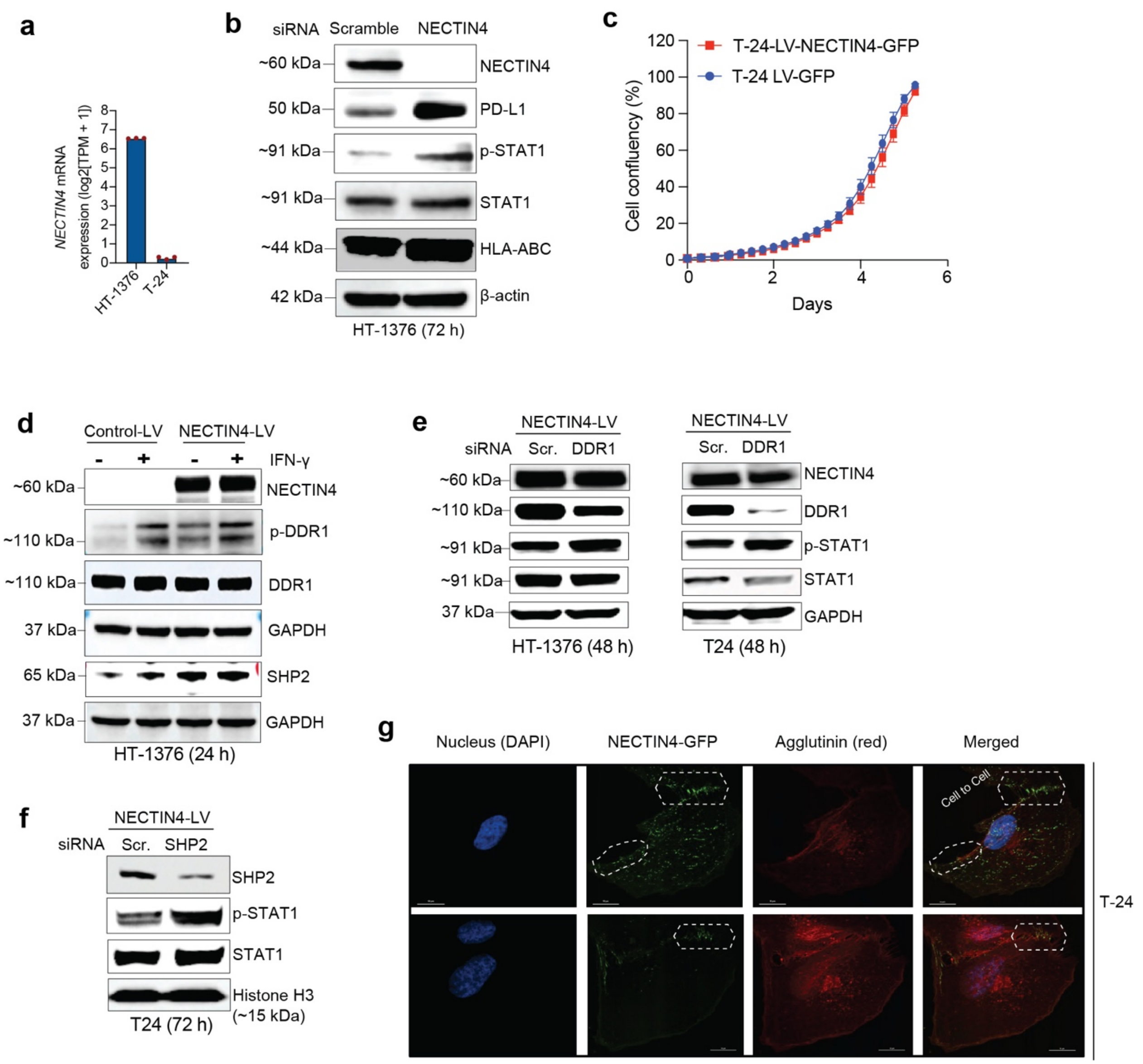


**Supplementary Fig. 8 | DDR1 or SHP2 knockdown abrogates NECTIN4-mediated STAT1 inhibition.**

**a,** Bar plot showing baseline mRNA expression of *NECTIN4* in HT-1376 and T24 cells, measured by RNA-seq.

**b,** Immunoblot analysis of NECTIN4, phospho (p)-STAT1 and total STAT1, HLA-ABC and PD-L1 in HT-1376 cells following siRNA-mediated NECTIN4 knockdown, with β-actin as a loading control.

**c,** Cell proliferation assays showing confluence over time for T24 cells expressing NECTIN4–GFP or control vectors, which lack endogenous *NECTIN4* expression. Confluence was quantified by live-cell imaging and normalized to baseline measurements.

**d,** Immunoblot analysis of phosphorylated DDR1, total DDR1 and SHP2 in HT-1376 cells expressing NECTIN4 or control vectors, in the presence or absence of IFN-γ (20 ng $ml^{-1}$).

**e,** Immunoblot analysis of HT-1376 and T24 NECTIN4-overexpression models following siRNA-mediated DDR1 knockdown, assessing NECTIN4, DDR1, phosphorylated STAT1 (p-STAT1) and

total STAT1. GAPDH was used as a loading control for HT-1376 cells, and histone H3 was used as a loading control for T24 cells.
**f,** Immunoblot analysis of T24 NECTIN4-overexpressing cells following siRNA-mediated SHP2 knockdown, assessing SHP2, phosphorylated STAT1 and total STAT1. Histone H3 was used as a loading control.
**g,** High-resolution confocal images showing DAPI (nuclei), NECTIN4–GFP and agglutinin (red). NECTIN4–GFP is enriched at cell–cell interfaces.

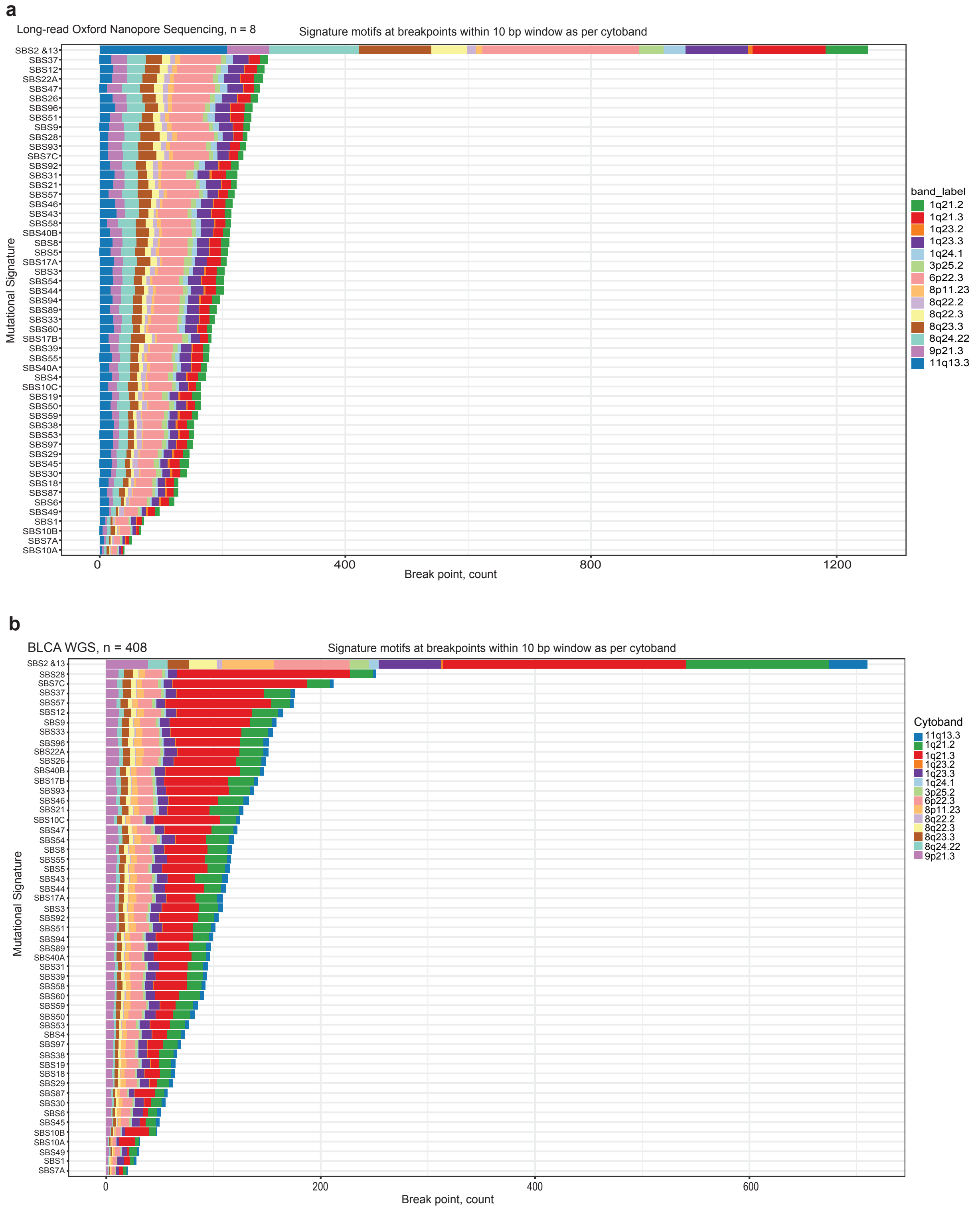


**Supplementary Fig. 9 | Whole-genome sequencing shows the distribution of mutational signature–specific motifs at breakpoints in copy-number–altered regions.**

**a,** Stacked horizontal bar plot showing the distribution of breakpoint-associated mutational signature motifs, stratified by cytoband, in long-read Oxford Nanopore whole-genome sequencing data from bladder tumors (*n* = 8).

**b,** Stacked horizontal bar plot showing the corresponding distribution of breakpoint-associated mutational signature motifs, stratified by cytoband, in TCGA bladder cancer whole-genome sequencing data (*n* = 408). Colors denote cytoband identity as indicated in the legend. Across both datasets, APOBEC3-associated mutational signatures (SBS2 and SBS13) account for the largest fraction of breakpoint motifs, with prominent enrichment at the chromosome 1q23 cytoband, which harbors recurrent *NECTIN4* amplifications.

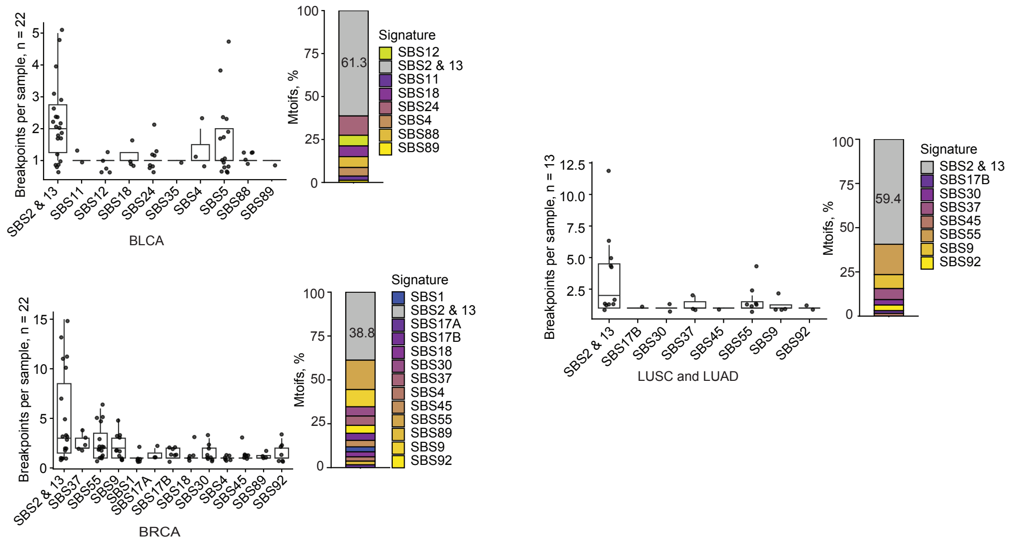


**Supplementary Fig. 10 | Signature-specific breakpoint motif coverage across *NECTIN4*-containing amplicons on chromosome 1q detected by AmpliconArchitect.**
Distribution of mutational signature–specific breakpoint motif coverage across chromosome 1q arm amplicons encompassing *NECTIN4* in BLCA, BRCA, LUAD, and LUSC tumors. Breakpoints were derived from GISTIC-defined *NECTIN4* amplification–positive samples in which AmpliconArchitect detected *NECTIN4* at ≥5 copies using default settings, enabling the capture of high-confidence breakpoints with sufficient read coverage. Each point represents an individual tumor sample. Stacked bar plots show the relative contribution of mutational signatures, highlighting the predominance of APOBEC-associated signatures (SBS2 and SBS13).
