## Supplementary Data Figures for "APOBEC3-driven neoantigen-rich cancers co-opt 1q23.3 amplification for tumor-intrinsic immune cloaking"

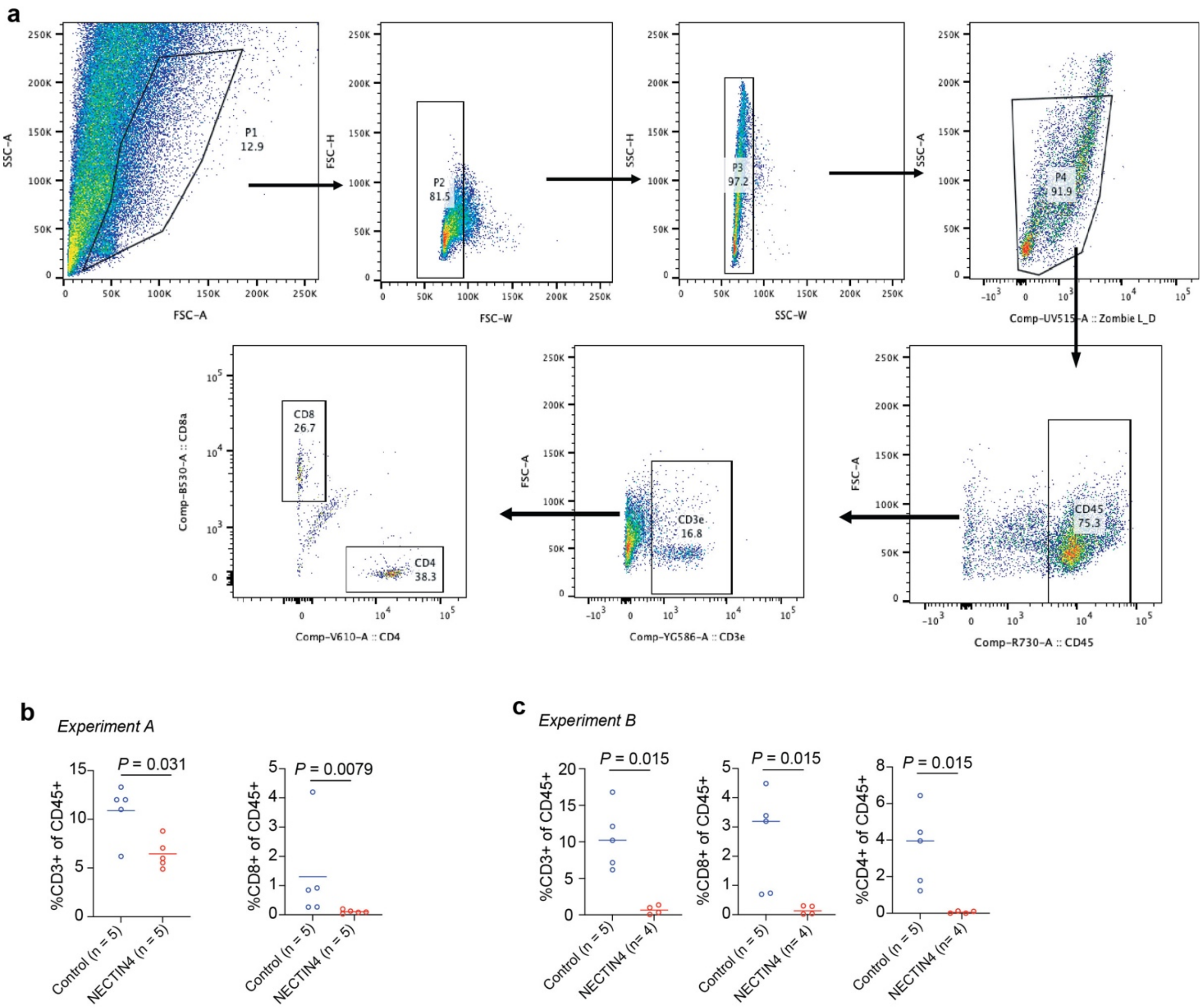


**Supplementary Data Fig. 1 | Flow-cytometry gating strategy and independent analysis of intratumoral T cells in NECTIN4-overexpressing tumors. a**, Representative flow-cytometry gating strategy used to quantify tumor-infiltrating T cells from syngeneic MB49 tumors implanted into immunocompetent C57BL/6 mice. Sequential gates were applied to exclude debris and doublets, followed by selection of live $CD45^+$ leukocytes and $CD3^+$ T cells. $CD8^+$ T cells were identified within the $CD3^+$ population. CD4 was included as an additional marker in Experiment B. b, Experiment A. Percentages of $CD3^+$ and $CD8^+$ T cells among $CD45^+$ tumor-infiltrating leukocytes in control and NECTIN4-overexpressing tumors. **c**, Experiment B. Percentages of $CD3^+$, $CD8^+$ and $CD4^+$ T cells among $CD45^+$ tumor-infiltrating leukocytes in control and NECTIN4-overexpressing tumors. Each dot represents an individual mouse horizontal lines indicate medians. P values were determined using two-sided non-parametric Mann–Whitney U tests.

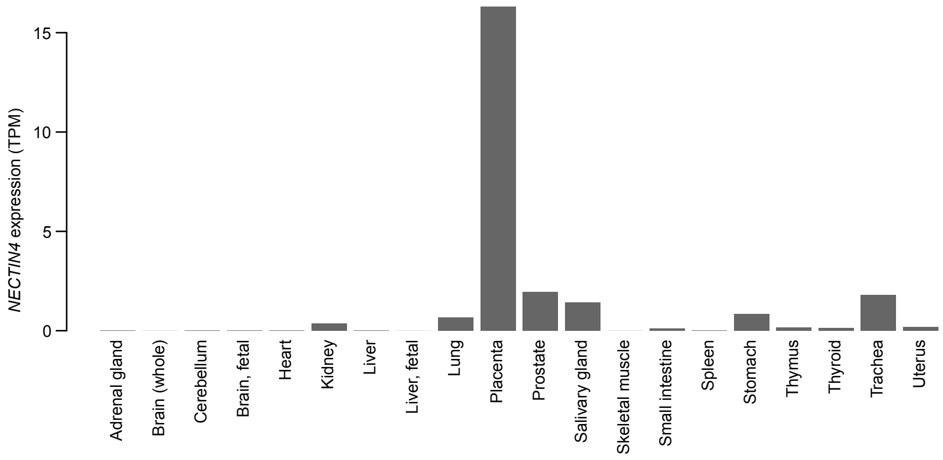


**Supplementary Data Fig. 2 | Expression pattern of *NECTIN4* across normal human tissues.** Bar plot showing mRNA expression levels of *NECTIN4* across normal human tissues based on RNA-seq data from BioProject PRJNA280600. Expression is highest in placenta.

BPS Bioscience # 78712

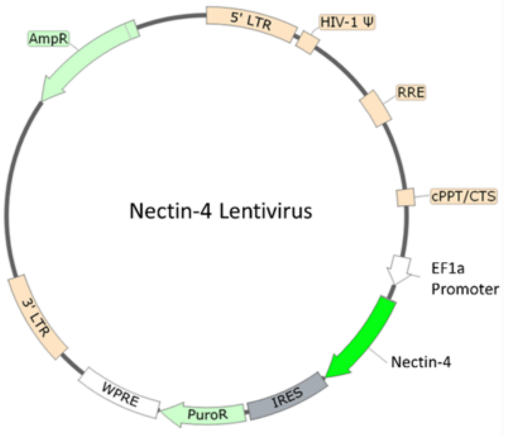


BPS Bioscience # 82212-P

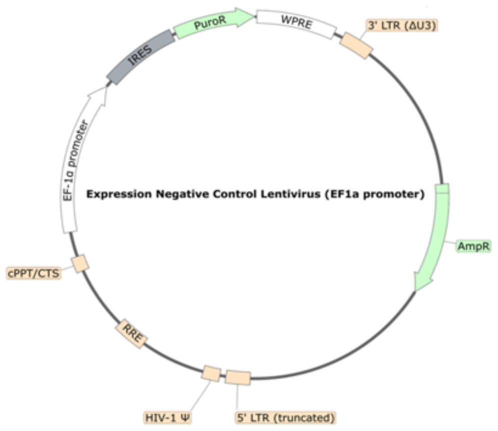


### Sequence Human Nectin-4 sequence (accession number: NM_030916.2)

MPLSLGAEMWGPEAWLLLLLLLLASFTGRCPAGELETSDVVTVVLGQDAKLPCFYRGDSGEQVGQVAWARVDAGEGAQELALL
HSKYGLHVSPAYEGRVEQPPPPRNPLDGSVLLRNAVQADEGEYECRVSTFPAGSFQARLRLRVLVPPLPSLNPGPALEEGQGLTLA
ASCTAEGSPAPSVTWDTEVKGTTSSRSFKHSRSAAVTSEFHLVPSRSMNGQPLTCVVSHPGLLQDQRITHILHVSFLAEASVRGLE
DQNLWHIGREGAMLKCLSEGQPPPSYNWTRLDGPLPSGVRVDGDTLGFPPLTTEHSGIYVCHVSNEFSSRDSQVTVDVLDPQE
DSGKQVDLVSASVVVVGVIAALLFCLLVVVVVLMSRYHRRKAQQMTQKYEEELTLTRENSIRRLHSHHTDPRSQPEESVGLRAEG
HPDSLKDNSSCSVMSEEPEGRSYSTLTTVREIETQTELLSPGSGRAEEEEDQDEGIKQAMNHFVQENGTLRAKPTGNGIYINGRG
HLV

Origene # RC203431L4

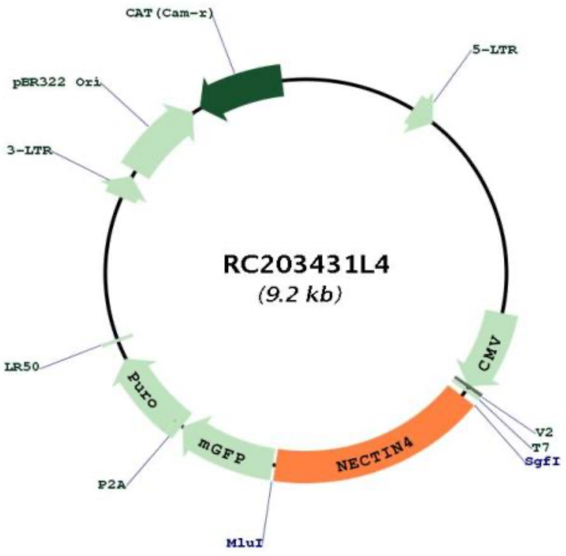


Origene # PS100071

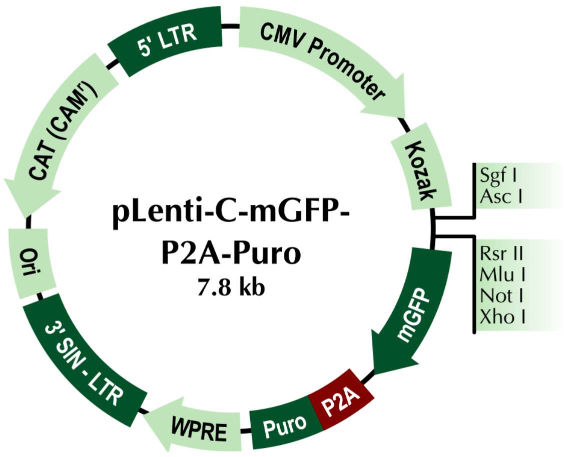

**Supplementary Data Fig. 3 | Plasmid constructs used in this study.** Schematic maps of lentiviral and expression vectors used for NECTIN4 overexpression and control experiments. Top left, EF1α-driven human NECTIN4 lentiviral construct (BPS Bioscience #78712). Top right, EF1α-driven negative control lentiviral vector (BPS Bioscience #82212-P). Bottom left, Origene RC203431L4 vector encoding full-length human NECTIN4 (NM_030916.2). Bottom right, pLenti-C-mGFP-P2A-Puro backbone (Origene PS100071) used for cloning and stable expression. The full-length human NECTIN4 coding sequence (accession number NM_030916.2) is shown below. All constructs were validated by sequencing prior to use.
